# Epigenetic Inheritance is Gated by Naïve Pluripotency and *Dppa2*

**DOI:** 10.1101/2021.05.11.443595

**Authors:** Valentina Carlini, Cristina Policarpi, Jamie A. Hackett

## Abstract

Environmental factors can trigger cellular responses that propagate across mitosis or even generations. Perturbations to the epigenome could underpin such acquired changes, however, the extent and contexts in which modified chromatin states confer heritable memory in mammals is unclear. Here we exploit a modular epigenetic editing strategy to establish *de novo* heterochromatin domains (epialleles) at endogenous loci and track their inheritance in a developmental model. We find that naïve pluripotent phases systematically erase ectopic domains of heterochromatin via active mechanisms, which acts as an intergenerational safeguard against transmission of epialleles. Upon lineage specification however, acquired chromatin states can be probabilistically inherited under selectively favourable conditions, including propagation of *p53* silencing through *in vivo* development. Using genome-wide CRISPR screening, we identify the mechanisms that block heritable silencing memory in pluripotent cells, and demonstrate removal of *Dppa2* unlocks the potential for epigenetic inheritance uncoupled from DNA sequence. Our study outlines a mechanistic basis for how epigenetic inheritance is restricted in mammals, and reveals genomic- and developmental-contexts in which heritable memory is feasible.

## INTRODUNCTION

Cellular identity is maintained by the constellation of trans- and cis-acting factors that regulate gene expression programmes. Amongst these, epigenetic mechanisms including histone modifications and DNA methylation, play a key role in establishing and perpetuating transcription states during development (Atlasi & Stunnenberg, 2017; Grosswendt *et al,* 2020). For example, heterochromatin domains facilitate stable transcriptional silencing, and are characterised by repressive H3K9me3 and DNA methylation or by H3K27me3 (Allshire & Madhani, 2018). Once established, DNA methylation patterns propagate through cell divisions via the maintenance methylase DNMT1, whilst histone modifications such as H3K27me3 and H3K9me3, utilise self-reinforcing feedback loops (Reinberg & Vales, 2018; Smith & Meissner, 2013). These entail ‘read-write’ modules that associate to the replication fork to reinstate modification patterns, and mutual cross-talk between epigenetic systems, which together are thought to promote stable ‘epigenetic’ inheritance. Nevertheless, chromatin marks are also subject to active reversal mechanisms and imperfect maintenance during replication, and are consequently rendered in a dynamic equilibrium of opposing influences. Thus, whilst chromatin states can convey a degree of heritable memory through reinforcing loops, they also exhibit plasticity in response to extrinsic cues.

These dual properties have implicated epigenetic systems as potential mechanisms that underlie genome-environment interactions (Cavalli & Heard, 2019). Indeed, across phyla and model organisms, environmental changes can induce specific epigenetic alterations – known as epialleles – that drive major phenotypic responses and adaptations (Duempelmann *et al*, 2019; Ge *et al*, 2018; Jiang & Berger, 2017; Seong *et al,* 2011; Simola *et al,* 2016; Torres-Garcia *et al,* 2020; Yang *et al,* 2017). In mammals, emergent phenotypes have also been linked with chromatin changes as a response to environmental contexts, for example hypoxia (Batie *et al,* 2019; Chakraborty *et al,* 2019), or availability of metabolic intermediates (Haws *et al,* 2020). Chromatin perturbations more generally are additionally associated with human disease susceptibility (Feinberg, 2018; Panzeri & Pospisilik, 2018). Understanding the potential prevalence and heritability of epialleles (or epimutations) in mammals is therefore of great interest.

Early embryogenesis is considered a susceptibility window for induction of epialleles (Bertozzi & Ferguson-Smith, 2020; Cavalli & Heard, 2019). Importantly, if epigenetic perturbations occur during development they have the potential to be inherited throughout adult tissues, possibly influencing disease risk (Hitchins, 2015; Walker, 2012). Furthermore, evidence is accruing across model organisms that adverse environments even prior to conception can induce epigenetic perturbations that are intergenerationally inherited, and influence offspring phenotype (Carone *et al,* 2010; Ciabrelli *et al,* 2017; Huypens *et al,* 2016; Klosin *et al,* 2017; Ost *et al,* 2014; Rechavi *et al,* 2014). However in mammals, preimplantation development entails reprogramming of parentally-inherited epigenomes, including rewiring of chromatin and global DNA demethylation (Hackett & Surani, 2013). Whilst this reprogramming is often believed to be directly linked with – or even prerequisite for – emergence of naive pluripotency, epigenome reprogramming could alternatively function principally as a barrier to inheritance of acquired or ectopic chromatin states (epialleles) (Kazachenka *et al,* 2018). In any case, the potential for heritable epialleles in mammals, and the underlying mechanisms that enable or antagonise this during development, are relatively uncharacterised.

The advent of epigenome editing tools has provided a means to program precise epigenetic perturbations at regulatory loci that can model environmentally-induced epialleles. Previous reports have suggested that co-targeted H3K9me3 and DNA methylation can exhibit stable propagation (Amabile *et al,* 2016; Bintu *et al,* 2016; Hathaway *et al,* 2012; Nunez *et al,* 2021), however these studies involve epigenetically abnormal cancer cell lines, do not extend *in vivo*, and/or manipulate underlying genetic contexts, whilst other studies found contradictory results (Braun *et al,* 2017; Kungulovski *et al,* 2015; Policarpi *et al,* 2021). Thus, the potential for epigenetic inheritance at endogenous loci in normal developmental contexts is unresolved. By exploiting an optimised and releasable CRISPR-dCas9 epigenetic editing tool to deposit broad heterochromatin domains, we reveal that developmental phases of naïve pluripotency function as a blockade to heritable silencing memory in mammals. Coupling our epigenetic memory assay with genome-wide genetic screens, we pinpoint *Dppa2* as a key ‘surveyor’ that restricts inheritance of epialleles in naïve cells. During subsequent developmental transitions however, inheritance of induced epialleles is permitted both *in vitro* and *in vivo*, but only under selective pressures. The results reveal heritable memory of induced chromatin states is viable in differentiated contexts, which has implications for disease risk, but places naïve pluripotency and *Dppa2* as an intergenerational safeguard against epiallele propagation in mammals.

## RESULTS

### A dynamic traceable system to programme de novo epialleles

To investigate the potential for memory of *de novo* epigenetic states we first developed an optimised CRISPR-based epigenetic programming tool. Here, we employed a catalytically-dead (d)Cas9 fused with an array of five optimally-spaced GCN4 repeats (dCas9^GCN4^) (Morita *et al,* 2016). These serve as docking sites to recruit up to five ‘effectors’ to a specific genomic locus via their single-chain antibody (scFv) domain (Fig 1A). This modular system amplifies the quantitative level and size of genomic domain of ON-target epigenome editing, relative to dCas9-effector fusions, whilst minimising OFF-target effects (Pflueger *et al,* 2018). To target *de novo* heterochromatin, we generated KRAB^GFP-scFv^ and DNMT3A/3L^GFP-scFv^ effectors, which promote direct deposition of H3K9me3 and DNA methylation (Quenneville *et al,* 2012).

**Figure 1.**
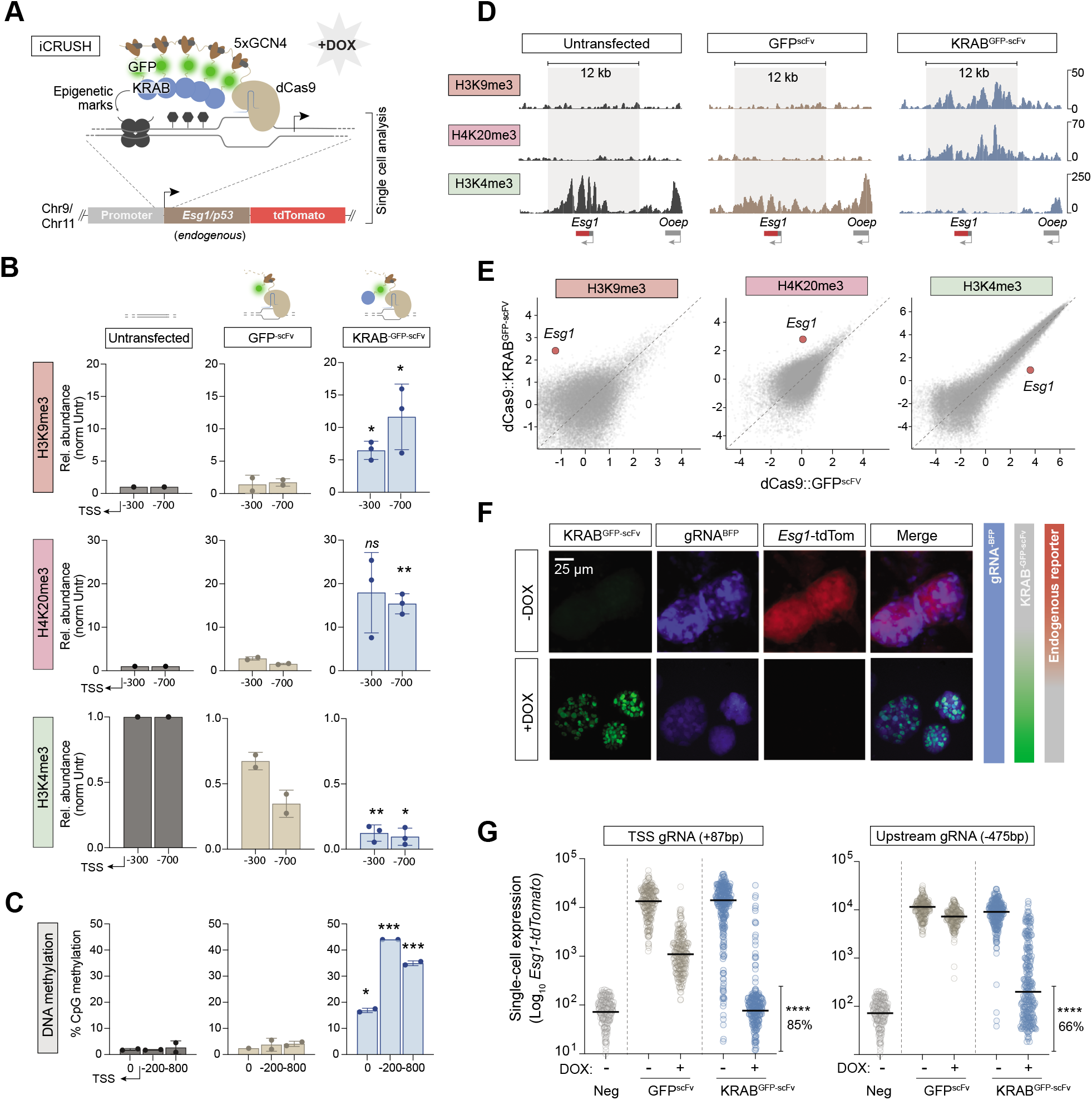
Programming epialleles with iCRUSH promotes robust heterochromatin domains and gene silencing. (**A**) Schematic showing recruitment of the modular epigenetic editing system to an endogenous promoter. The inducible CRISPR unleashing of silencing by heterochromatin (iCRUSH) tool is DOX-inducible and destabilsed enabling dyamic temporal control, and carries GFP for real-time tracking. It recruits five KRAB^GFP-scFV^ modules via a GCN4 array to promote enhanced ON-target epigenetic editing over large chromatin domains. A knock-in reporter downstream of target loci facilitates single-cell analysis. (**B**) The relative abundance of indicated histone modifications assayed by CUT&RUN-qPCR after KRAB^GFP-scFv^ (blue) or control GFP^scFv^ (light brown) targeting, relative to untransfected (grey). Shown are independent quantifications at two genomic positions on the *Esg1* promoter, indicated relative to TSS (−300, −700). Data is average of two or three biological replicates. (**C**) Histograms showing the average DNA methylation across three genomics regions of the *Esg1* promoter determined by bisulfite pyrosequencing. (**D**) CUT&RUN genome tracks for H3K4me3, H4K20me3, and H3K4me3 in untransfected (grey), control GFP^scFv^ (light brown) or KRAB^GFP-scFv^ (blue) targeted ESC after 7 days of DOX treatment. Grey box highlights the region of heterochromatin spreading induced by epigenetic editing. (**E**) Scatterplots demonstrating specificity and magnitude of programmed modifications at *Esg1* by CUT&RUN-seq. Shown are all promoters genome-wide. (**F**) Epifluorescence images of *Esg1^tdTomato^* ESCs in -DOX (top) or +DOX conditions (bottom), where targeted heterochromatin is deposited. (**G**) Single-cell expression of *Esg1^tdTomato^* in control (GFP^-scFv^) or or upon heterochromatin induction (KRAB^-GFP-scFv^) using a gRNA targeting close to the TSS (+87bp) or further upstream on the promoter (−475bp). Each datapoint indicates a cell, percentage indicates the proportion fully silenced cells (mean of four biological replicates), bars represent median. In all panels, asterisks indicate *p-values* by unpaired t-test; *\*p<0.05, **p<0.01 ***p<0.001.Error* bars ± S.D.

We placed all components under a DOX-inducible promoter and destabilised dCAS9^GCN4^ protein and effectors with d2 domains, which together facilitate precise temporal control over epigenome editing activity. This is important to assess subsequent epigenetic memory without confounding reiterative targeting. To track the temporal ON-OFF dynamics in real-time, all effectors are tagged with superfolder-GFP, which also enables cell isolation via flow cytometry (Fig S1A-C). Finally, we utilised an ‘enhanced’ gRNA scaffold (AT-flip, extended stem loop) linked with a tagBFP (Chen *et al,* 2013), which further amplifies ON-target activity and facilitates tracking, respectively. In summary, dCas9^GCN4^ and KRAB^GFP-scFv^ expression can be induced by DOX treatment and traced in real-time via GFP, while BFP is constitutively expressed. Reciprocally, the system is destabilised and can be rapidly switched back OFF by removal of DOX.

### Programmed heterochromatin epialleles fully silence gene activity in single-cells

To follow programmed epialleles at endogenous loci, we initially used a naïve pluripotent ESC line wherein the endogenous *Esg1* gene carries a knock-in tdTomato (Fig 1A) (Hackett *et al,* 2018). We introduced dCas9^GCN4^::KRAB^GFP-scFV^ and a single gRNA^BFP^ that targets the *Esg1* promoter via piggyBac, and assessed the extent of *de novo* programmed epigenetic states after 7 days (7d) induction with DOX. Quantitative CUT&RUN-qPCR demonstrated a highly significant deposition of heterochromatic H3K9me3 (*p=0.014*) marks across the *Esg1* promoter specifically with KRAB^GFP-ScFv^, relative to either untransfected or control GFP^ScFv^ targeting (Fig 1B). This was paralleled by enrichment of another heterochromatic mark, H4K20me3 (*p=0.002*), which often colocalises with H3K9me3 (Schotta *et al,* 2004), and complete loss of endogenous H3K4me3 modification (*p=0.0013*). Moreover, bisulfite pyrosequencing revealed a highly significant increase of DNA methylation (*p=0.0005*) across the entire *Esg1* promoter region (Fig 1C). These results indicate that upon single-gRNA tethering of a flexible array of five KRAB^GFP-scFV^ effectors, a *de novo* domain of heterochromatic modifications is established.

To further investigate the extent and specificity of programmed heterochromatin we performed CUT&RUN-seq. We observed that our epigenetic editing system deposits a broad domain encompassing ~12kb of H3K9me3 and H4K20me3 around the endogenous *Esg1* locus, whilst previously abundant H3K4me3 is undetectable (Fig 1D). Importantly the *de novo* peaks of H3K9me3 and H4K20me3 are of a magnitude comparable to the strongest peaks throughout the genome, suggesting they recapitulate robust physiological heterochromatin status (Fig 1E & S1D). Moreover, targeting was highly specific, since we observed minimal OFF-target changes in H3K9me3, H4K20me3 and H3K4me3 (Fig 1E & S1D). Taken together, these data reveal that a substantial epigenomic domain (>10kb), which bears the key hallmarks of repressive heterochromatin, is specifically programmed at an endogenous genomic locus.

We next asked whether this *de novo* heterochromatin domain is associated with induction of transcriptional silencing. *Esg1* is highly active in pluripotent cells, and the endogenous reporter facilitates dynamic single-cell analysis over time. As expected, addition of DOX led most cells to become GFP-positive, indicative of activation the epigenetic editing system (Fig 1F). Strikingly, this concomitantly led to complete loss of *Esg1*^tdTomato^ positive cells, which was additive with time (Fig 1F & S1C). Quantitative single-cell expression revealed >99% cells exhibited transcriptional repression, with >85% reaching a complete OFF state, indistinguishable from ESC that do not carry the tdTomato reporter (Neg), and corresponding to >500-fold transcriptional silencing (Fig 1G). In contrast the GFP^scFV^ control exhibited only modest repression, which likely reflected steric hindrance due to transcriptional start site (TSS) binding, since a gRNA targeting upstream elicited full silencing with KRAB^GFP-scFV^ (>500 fold) but its control GFP^scFV^ exhibited no effect on transcription (Fig 1G). In summary, these data indicate that our system is able to ectopically programme heterochromatin states at *Esg1*, which is directly linked with powerful transcriptional silencing at the single-cell level, implying high penetrance of deposition across a broad domain. We refer to this enhanced epigenetic tool as <u>i</u>nducible <u>CRI</u>SPR <u>u</u>nleashing of <u>s</u>ilencing by <u>h</u>eterochromatin (iCRUSH) (Fig 1A).

### Induced heterochromatin is progressively erased in ESC

Extant paradigms suggest that large domains of heterochromatic H3K9me3, H4K20me3 and DNA methylation are heritable, and self-propagate via ‘read-write’ feedback machinery (Hathaway *et al.*, 2012; Reinberg & Vales, 2018). To understand this in a developmental context, we next investigated the potential for propagation of induced heterochromatin epialleles in naïve ESC, which faithfully recapitulate *in vivo* epiblast when maintained in 2i/L (Hackett & Surani, 2014). Withdrawal of DOX resulted in a rapid switch off of the iCRUSH epigenetic editing system as determined by quantitative cytometry for GFP, and consistent with dCas9^GCN4^ and KRAB^GFP-scFV^ being destabilised, therefore fully releasing the inducing signal (Fig 2A). In parallel we observed a progressive loss of *Esg1*^tdTomato^ silencing (Fig 2B). Interestingly, 4d after DOX washout (D-wo (4d)) we observed a bimodal distribution of *Esg1* expression amongst single-cells, indicative of probabilistic reactivation dynamics. By 7d after release (D-wo (7d)) however, all cells reverted to the ON state, reflecting >500-fold increase in *Esg1* expression (Fig 2B).

**Figure 2.**
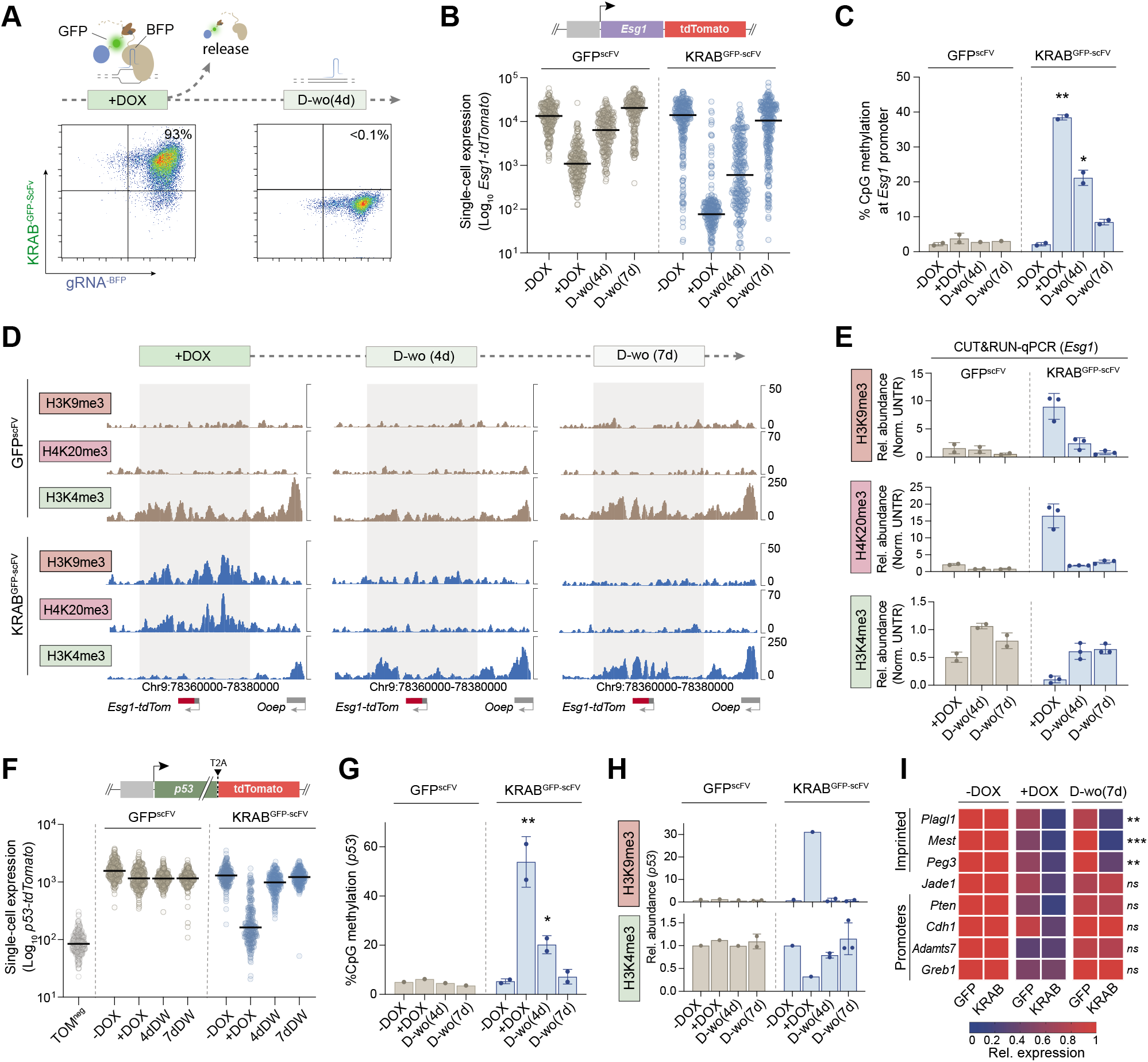
Induced heterochromatin epialleles are progressively erased in naïve pluripotent cells. (**A**) Representative flow-cytometry density plots showing gRNA^BFP^ and KRAB^GFP-scFv^ expression after DOX treatment (7days) and DOX washout (4 days). (**B**) *Esg1^tdTomato^* expression at single-cell resolution during DOX washout in control (GFP^-scFv^) or induced heterochromatin (KRAB^-GFP-scFv^) cells. Horizontal bars indicate median, each datapoint a single cell. (**C**) Histograms of mean DNA methylation levels across the *Esg1* promoter (6 CpG sites) during DOX washout in independent experiments. Statistics are measured comparing individual timepoints with the -DOX control. (**D**) CUT&RUN tracks at +DOX, and 4 and 7 days of DOX washout in control GFP^scFv^ or KRAB^GFP-scFv^ for indicated histone marks. Grey boxes highlight the domain of heterochromatin spreading in +DOX. (**E**) CUT&RUN-qPCR quantification of the relative abundance of each mark at *Esg1* promoter in independent biological replicates, normalised to a positive control region and untransfected cells. (**F**) *p53*^tdTomato^ expression in single-cells during DOX washout in control (GFP^-scFv^) or induced heterochromatin (KRAB^-GFP-scFv^) cells. (**G**) Bisulfite pyrosequencing quantification of DNA methylation at the *p53*-promoter (4 CpG sites) at indicated timepoint. (**H**) CUT&RUN-qPCR quantification of the relative abundance of H3K9me3 and H3K4me3 at *p53* promoter in independent biological replicates. (**I**) Heatmap representing relative expression by qPCR of each indicated gene upon heterochromatin targeting (+DOX) or after DOX washout (D-wo (7d)), normalized to the -DOX control in three biological replicates. In all panels, asterisks indicate *p-values* by unpaired t-test; *\*p<0.05, **p<0.01 ***p<0.001*. Error bars ± S.D.

To determine if transcriptional re-expression corresponds to loss of programmed epigenetic states we used bisulfite pyrosequencing and CUT&RUN. Consistently with reactivation of expression, we found that DNA methylation is partially maintained at the *Esg1* promoter at the early timepoint (D-wo (4d)) but is almost completely erased by 7 days (D-wo (7d)) (Fig 2C). However, we found that the high levels of deposited H3K9me3 and H4K20me3 are largely erased by 4d after DOX withdrawal (Fig 2D-E). Following 7d release of the epigenetic editing trigger, the *Esg1* chromatin state completely reverts to the initial configuration, including erasure of H3K9me3, H4K20me3 and DNA methylation, and reacquisition of the endogenous H3K4me3 mark (Fig 2D-E). Whilst our system deposits high levels of DNA methylation, we additionally checked whether co-targeting KRAB^GFP-scFv^ together with the catalytic domain of *Dnmt3a* and its cofactor *Dnmt3L* (3a3L^GFP-scFv^) would enhance epigenetic inheritance, since such effects have been reported in cancer cell lines (Fig. S2A) (Amabile *et al.*, 2016). Although we found a slight further increase of DNA methylation by compound recruitment (Fig S2B), we observed equivalent or faster erasure of epigenetic memory (Fig. S2C). Taken together, our data argue that induction of a robust domain of heterochromatin, and consequently extensive epigenetic silencing, is readily reversible from OFF→ON in naïve pluripotent cells.

### Context-dependent influences of epigenetic inheritance in ESC

To confirm that failure to propagate robust heterochromatin in ESC is not a phenotype specific to *Esg1*, we generated a second endogenous reporter ESC line by inserting *tdTomato* downstream of the *p53* gene, separated by a T2A self-cleavable domain (Fig. 2F). Targeting KRAB^GFP-scFv^ to the *p53*^tdTomato^ promoter recapitulated the same extensive heterochromatin deposition including DNA methylation, H3K9me3 and loss of H3K4me3, and robust single-cell silencing, as achieved at *Esg1*^tdTomato^ (Fig. 2F-H). Upon 7d DOX withdrawal however, we found that *p53* expression becomes fully reactivated in ESC (Fig. 2F-H). This is paralleled by erasure of the targeted DNA methylation and H3K9me3, and reacquisition of endogenous H3K4me3, consistent with heterochromatin failing to confer epigenetic memory in naïve ESC.

To further examine this principle across distinct genomic locations, we programmed heterochromatin to additional endogenous loci, selected to represent different regulatory features (*e.g*. imprinting control regions, promoters). We imposed strong epigenetic silencing with iCRUSH, yet most loci (*Pten, Cdh1, Greb1, Adamts7* and *Jade1*) reverted to their original expression status within 7d DOX withdrawal regardless of their initial epigenetic state, promoter features (CpG density) or initial expression level (Fig. 2I), consistent with *Esg1* and *p53* dynamics. Nevertheless, we did observe that imprinted genes (*Peg3, Mest* and *Plagl1*) are exceptions and, uniquely, maintain memory of *de novo* silencing in naïve ESC (Fig 2I). This suggests that heterochromatin domains at ectopic sites can be epigenetically inherited in a genomic context-dependent manner, with imprinted loci providing the necessary sequence substrate for heterochromatin propagation. However in general, we find *de novo* chromatin states at endogenous single-copy loci are not heritable over extended periods in naïve ESC. This suggests a dynamic competition of opposing activities that generally disfavours epigenetic inheritance during the phase of early developmental naïve pluripotency, potentially as a safeguard to restrict intergenerational transmission of aberrant epialleles.

We next asked whether the reversal of heterochromatic epialleles in ESCs is driven by passive dilution through cell divisions or by active erasure. For this, we used transient transfection of a gRNA to epigenetically silence the *Esg1* reporter for 3 days by DOX treatment. This led to ~100-fold silencing, deposition of significant levels of DNA methylation, H3K9me3 and H4K20me3, and loss of H3K4me3 (Fig S3A-B). We then released the epigenetic editing system by DOX washout and concomitantly treated the cells with or without the cell cycle inhibitor RO3306 (Fig S3C), which blocks cells at the G2/M phase boundary. We observed *Esg1* reactivation is only weakly impaired by cell cycle inhibition (Fig S3D), spanning at least 60 hours (Fig S3E). Thus, whilst passive dilution may partially contribute to reversion of epigenetic memory, active mechanisms play a key role in erasing *de novo* epialleles in ESC.

### CRISPR screen reveals key factors that antagonise epigenetic memory in ESC

We therefore sought to identify the putative factors that actively counteract epigenetic memory in pluripotent phases by designing a genome-wide loss-of-function CRISPR screen (Fig 3A). We introduced a single-copy of *Cas9* nuclease tagged with GFP (Cas9^T2A-GFP^) into *Esg1*^tdTomato^ ESC that also carry dCas9^GCN4^ in the OFF state, and infected these cells with a pooled lentiviral library of exon-targeting gRNA (Doench *et al,* 2016). We subsequently induced self-inactivation of the Cas9^T2A-GFP^ with a pair of specific gRNAs, which we confirmed by flow sorting cells according to loss of GFP (GFP^neg^) (Fig 3A). This cell population is now composed of a heterogeneous pool of knockout cells, to which we could apply our epigenetic editing system to identify the factors that antagonise epigenetic inheritance.

**Figure 3.**
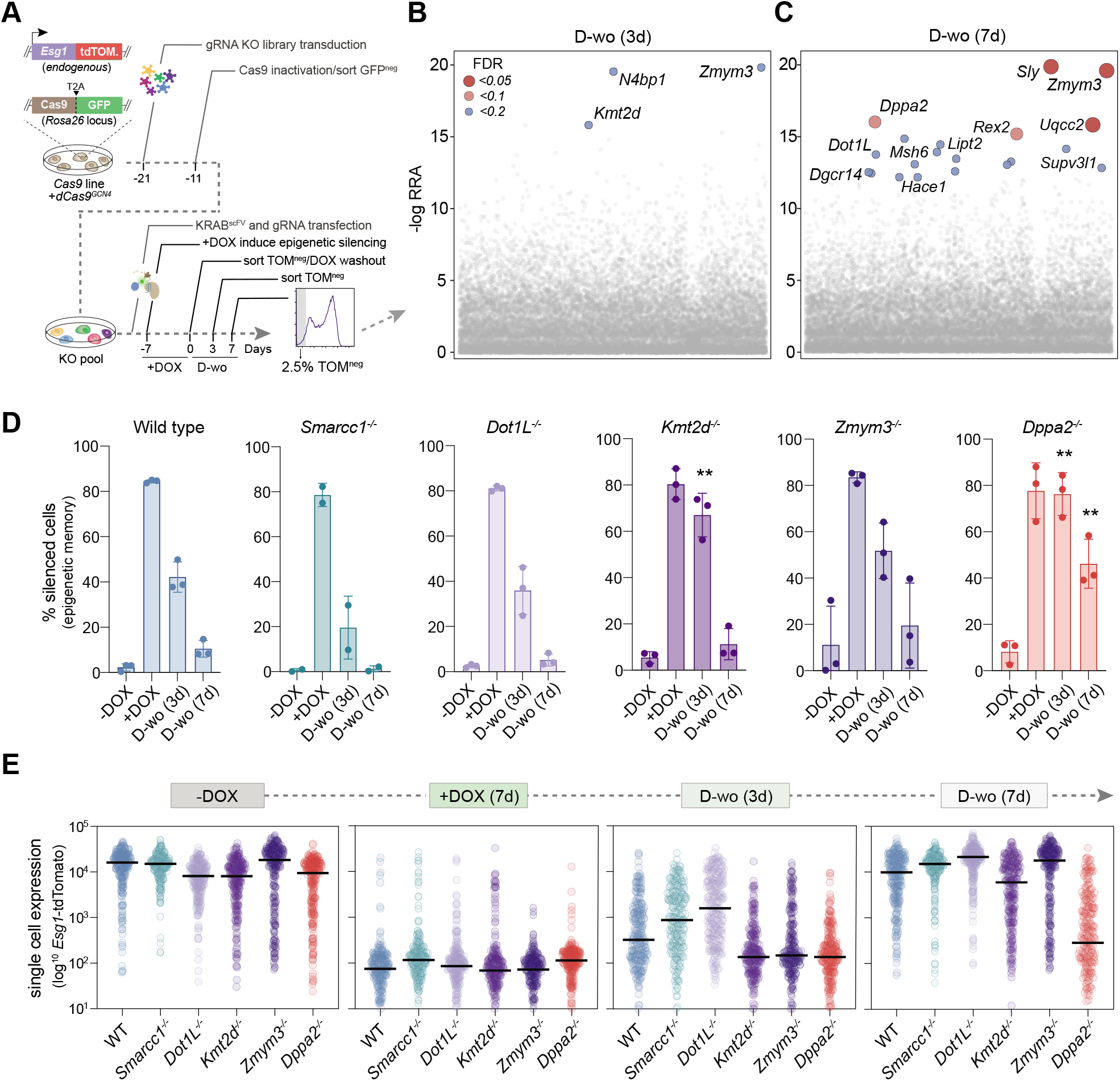
A CRISPR screen identifies key factors that restrict epigenetic inheritance in naive ESC. (**A**) Schematic of screen design and workflow: a *Esg1^-tdTomato^* cell line carrying constitutive *Cas9*-GFP nuclease and DOX-inducible (*off*)-*dCas9^GCN4^* is transduced with a lentiviral gRNA library. The *Cas9-*GFP is self-inactivated via introducing specific gRNAs, and GFP-negative cells are isolated. Subsequently, gRNA^BFP^ and KRAB^GFP-ScFv^ are introduced and cells treated with DOX to induce *Esg1* heterochromatin-mediated silencing (TOM^neg^). TOM^neg^ ESC are flow sorted and re-cultured in absence of DOX to isolate cells that inherit epigenetic silencing, which are subject to NGS to identify the gene knockout they carry. (**B-C**) Significant hits for factors that permit epigenetic memory when knocked-out, showing the -log relative ranking algorithm (RRA) score at 4 or 7 days of DOX-washout. False discovery rate (FDR) is indicated. (**D**) Histograms showing the percentage of *Esg^tdTomato^* negative cells in WT or independent knockout ESC after programming heterochromatin (+DOX) and during DOX-washout. Each datapoint indicates a biological independent knockout line. Error bars ± S.D, asterisks indicate *p-values* relative to -DOX by unpaired t-test; *\*p<0.05, **p<0.01.(**E***) Quantitative expression of *Esg^tdTomato^* amongst single-cells in WT or in independent clonal knockout ESC lines. Bars represent the median.

To achieve this, we targeted heterochromatin to *Esg1*^tdTomato^ and isolated cells that subsequently retained silencing memory (TOM^neg^) following release of dCas9^GCN4^::KRAB^GFP-scFv^ using a gating strategy to distinguish between cells remaining fully silenced (bottom 2.5% (TOM^neg-2.5%^) (Fig S4A) and those that reatain a degree of repression memory (TOM^neg-wide^) (Fig S4B). We then used model-based analysis of genome-wide CRISPR-Cas9 knockout (MAGeCK) to identify the gene knockouts enriched in the TOM^neg^ populations that retained epigenetic memory relative to the complementary TOM^pos^ population over short- (3d) and extended- (7d) timescales (Li *et al,* 2014). As expected, top hits across both gates were associated with roles in transcriptional or translational regulation, and comprised many candidates with established or predicted epigenetic functions. This included the SWI/SNF histone remodeller *Smarcc1* (FDR 0.03), the H3K79 methyltransferase *Dot1L* (FDR 0.16) and the H3K4 histone methyltransferase *Kmt2d* (FDR 0.13), although this latter was enriched only at the shorter timepoint (Fig 3B-C & S4B). Additionally, we noted cells that propagated silencing memory also exhibited significant enrichment for knockouts of the X-linked zinc-finger protein *Zmym3* (FDR 0.01), the NSL complex subunit *Kansl2* (FDR 0.05), and *Dppa2* (FDR 0.03) (Fig 3B-C, S4B-C), which was recently linked with regulating *de novo* DNA methylation and bivalency (Eckersley-Maslin *et al,* 2020; Gretarsson & Hackett, 2020).

To validate these candidates, we generated independent clonal knockout ESC lines of each. Deletion of these factors did not affect *Esg1*^tdTomato^ basal activity prior to imposition of epigenetic silencing, and all knockouts also exhibited a comparable extent of programmed silencing as WT after 7d DOX, implying no changes in initial parameters (Fig 3D). Following DOX-washout, *Smarcc1^-/-^* and *Dot1L^-/-^* reverted to the active state with a similar kinetics to WT, implying false-positives. In contrast, *Kmt2d*^-^ cells showed penetrant memory at 4d of DOX washout, but reverted to an ON state after 7 days, suggesting absence of *Kmt2d* impacts the rate of memory erasure, potentially by affecting redeposition of H3K4me3 (Fig 3D-E). Interestingly whilst some *Zmym3^-/-^* lines exhibited memory, and independent gRNAs were concordant (Fig S4C), there was high heterogeneity between independent knockout clones, indicating a complex regulatory response that we did not follow further. Remarkably however, all *Dppa2^-/-^* lines fully maintained epigenetic memory after 3d DOX-withdrawal, with the majority of the cells remaining in the OFF state after 7days (Fig 3D-E & S4C). This suggests that abrogating *Dppa2* changes the balance of activates in ESC to generate an environment that is permissive for epigenetic inheritance.

### Epigenetic inheritance is permitted by deletion of DPPA2

To examine the role of *Dppa2* further, we traced the single-cell dynamics of transcriptional memory in multiple knockout ESC lines (Fig S5A). Whilst WT cells rapidly lose silencing memory after 7days DOX washout, the majority of *Dppa2^-/-^* cells remain fully silenced at this stage (Fig. 4A). Importantly, inheritance of this silenced state in the absence of *Dppa2* is subsequently maintained across a consistent fraction of cells for at least 43 days after DOX withdrawal (>100 cell replications), with the population therefore exhibiting a bimodal distribution (Fig. 4B). Importantly doubling time was very similar between wildtype and knockout cells (Fig S5B). This implies that abrogation of *Dppa2* facilitates heritability of ectopic silencing in a probabilistic manner, potentially by shifting the odds against reversion, and thus promoting steady-state inheritance. Notably, flow sorting TOM^neg^ and TOM^pos^ fractions after 26 days of DOX washout (revealed that, while the TOM^pos^ remained positive, the TOM^neg^ reacquired a bimodal distribution, supporting a stochastic memory function (Fig S5C-E).

**Figure 4.**
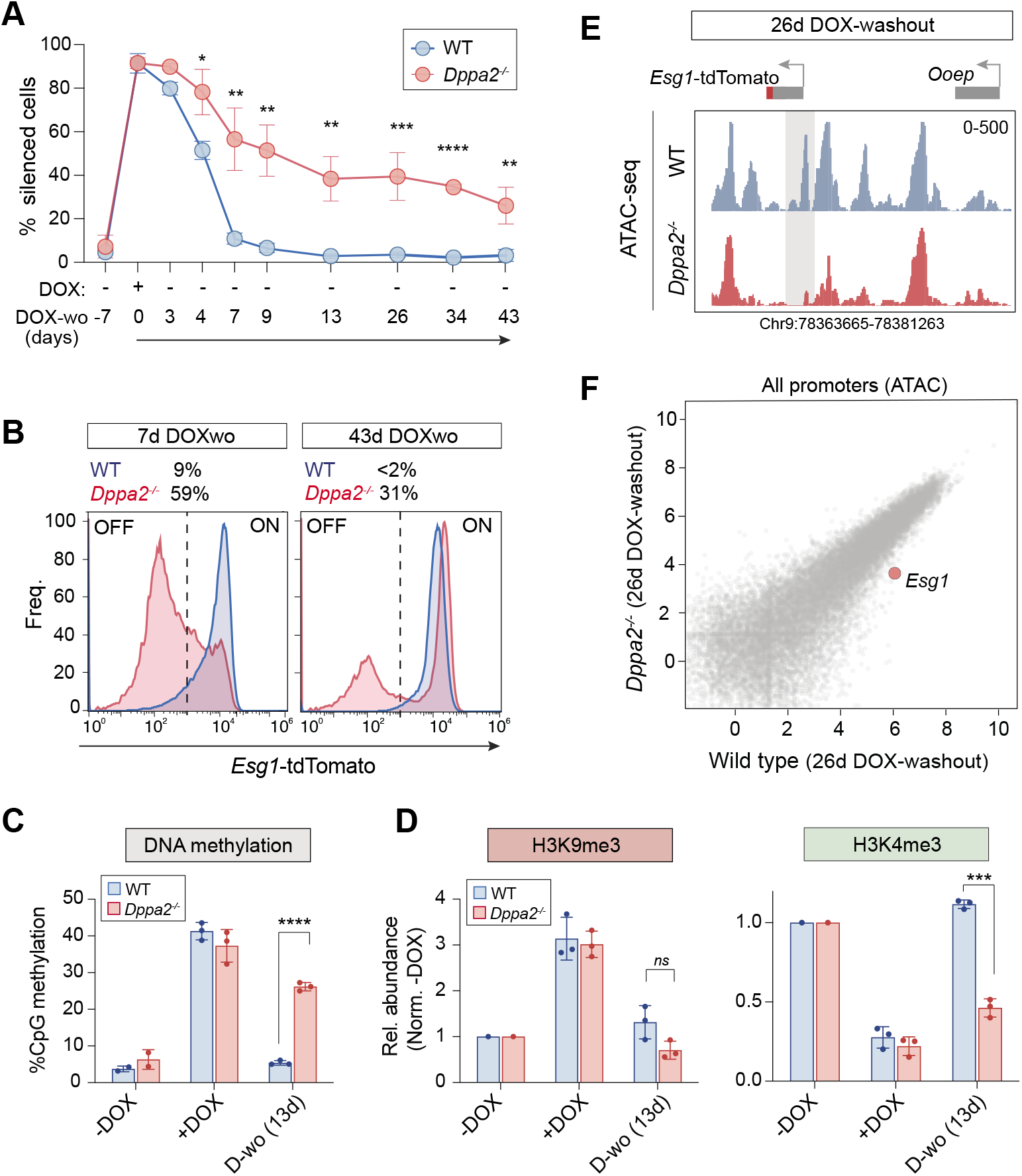
Epigenetic inheritance is unlocked by deletion of *Dppa2*. **(A)** Time-course showing percentage cells that propagate heritable epigenetic silencing of *Esg1^tdTomato^* in wildtype or *Dppa2^-/-^* cells. Each datapoint represents average of three biological independent lines. (**B**) Distribution of quantitative *Esg1^tdTomato^* expression in *Dppa2^-/-^* (red) or wildtype (blue) after 7 or 43 days DOX washout showing bimodal epigenetic memory only upon *Dppa2* abrogation. Numbers indicate percentage of *Esg1^tdTomato^*-negative cells. (**C**) Bisulfite pyrosequencing quantification of DNA methylation at *Esg1* promoter in WT (blue) or Dppa2^-/-^ (red) cells, each assayed in three independent lines. (**D**) CUT&RUN-qPCR quantification of H3K9me3 and H3K4me3 relative to a positive control region and to the -DOX control. (**E**) ATAC-seq tracks showing chromatin accessibility at *Esg1^-tdTomato^* after 26 days of DOX washout in wildtype or *Dppa2^-/-^* lines. Grey box highlights *Esg1* promoter. (**F**) Scatterplot of genome accessibility across all promoters after 26d DOX-withdrawal, showing a specific and persistent memory at *Esg1* in *Dppa2^-/-^* cells. In all panels, asterisks indicate *p-values* by unpaired t-test; *\*p<0.05, **p<0.01 ***p<0.001*. Error bars ± S.D.

We next investigated transmission of induced epigenetic states in *Dppa2^-/-^* cells. After DOX induction of iCRUSH, we observed that DNA methylation and H3K9me3 are deposited comparably in both WT and *Dppa2^-/-^* cells, and endogenous H3K4me3 is equivalently erased (Fig 4C-D). Upon release of dCas9^GCN4^::KRAB^GFP-scFv^ however (DOX washout), whilst WT cells underwent a complete recovery of the epigenetic landscape, *Dppa2^-/-^* exhibited highly significant inheritance of DNA methylation (Fig 4C), and also propagated the H3K4me3 depleted state (Fig. 4D). Moreover, ATAC-seq revealed that induced chromatin inaccessibility status was transmitted mitotically in *Dppa2^-/-^* ESC but not WT (Fig. 4E). This reflected a specific response at epigenetically perturbed loci, since non-targeted promoters did not exhibit accessibility changes in the absence of *Dppa2* (Fig. 4F). To determine the generality of epiallele propagation in *Dppa2^-/-^* ESC, we targeted heterochromatic silencing to additional loci, which we previously showed do not exhibit memory in wildtype ESC (Fig 2I). We observed propagation of repression at *Pten, Cdh1, Greb1 and Adamts7* specifically in the absence of *Dppa2* (Fig. S5F). Whilst this did not reach significance, this likely reflects an incompletely penetrant memory at the single cell level, similar to *Esg1*. Nonetheless, we did observe consistantly significant inheritance of an induced repressed state at *Jade1* in *Dppa2^-/-^* compared to wild-type (Fig. S5F).

Taken together these data suggest that once an aberrant heterochromatic state occurs, pluripotent cells rely, at least partially, on DPPA2 to re-establish the original epigenetic configuration. This implies DPPA2 acts as an epigenome ‘surveyor’ in naïve pluripotent cells by promoting probabilistic reversion of epimutations.

### Aberrant epialleles can be propagated upon exit from pluripotency

Our data indicate that naïve ESC robustly reverse ectopic epigenetic states at endogenous loci, which hinges on *Dppa2*. Notably, *Dppa2* is only expressed during pluripotent phases suggesting epialleles acquired during or after this may confer heritable mitotic memory through subsequent lineage commitment. To investigate this, we differentiated wild-type ESC towards definitive endoderm as an *in vitro* model of development (Fig. 5A). Because *Esg1* is repressed during differentiation as part of the normal developmental programme, we initially focused on *p53*, wherein heterochromatin and transcriptional silencing are also robustly erased in ESC (Fig 2F-H).

In contrast to ESC, upon differentiation to definitive endoderm we observed a highly penetrant memory of *p53* silencing amongst single-cells (Fig. 5B), whereas no memory effects were observed upon control targeting with GFP^scFV^ (Fig S6A). Analysis of chromatin revealed full inheritance of targeted DNA methylation (>85%) at *p53* specifically in differentiating endoderm cells (Fig 5C), whilst there is also heritable memory of the H3K4me3 depletion (Fig 5D). Interestingly, deposited H3K9me3 is erased in endoderm (Fig 5D), implying it does not self-reinforce nor drive silencing in this context.

To investigate this epigenetic memory further, we used ATAC-seq and observed highly significant loss of accessibility specifically at targeted *p53* upon *de novo* heterochromatin formation (Fig 5E & S6B). Following 7d DOX withdrawal this inaccessible chromatin state exhibited robust memory during endoderm differentiation. In contrast, chromatin accessibility is restored in ESC upon DOX withdrawal (Fig 5E). These data imply that differentiated cells, but not naïve pluripotent ESC, are competent for epigenetic inheritance of ectopic heterochromatin.

### Epigenetic inheritance is probabilistic and highly context-dependent

To further understand the principles that underpin epigenetic inheritance in specific contexts, we reasoned that *p53* epigenetic silencing may facilitate a selective advantage. To test this, we performed a growth assay by mixing equal (1:1) proportions of silenced *p53* cells with untargeted *p53* controls carrying constitutive GFP expression (CAG:GFP). In ESC, we observed no selective advantage of prior *p53* silencing following DOX washout (GFP^neg^), possibly due to rapid reversion mechanisms in ESC overcoming any growth advantage. In contrast upon endoderm induction, cells that were previously targeted for *p53* epigenetic silencing quickly become dominant in the population, comprising >95% by d8 following co-culture (Fig. 5F). This implies that initial *p53* silencing confers a strong advantage in differentiating cells, even after the inducing trigger is removed, suggesting any erasure mechanisms are outcompeted by the growth advantage. Indeed, we observed prior *p53*-silenced endoderm replicate faster with greater viability (Fig. 5G).

Because erasure of heterochromatin at *Esg1* and *p53* occurs probabilistically after DOX-withdrawal (generating a transient bimodal distribution), we propose that the balance of opposing factors that influence epigenetic inheritance or erasure is shifted by the growth advantage conferred by epigenetic silencing at *p53*. In this scenario, there is an interaction between genetics and epigenetics that facilitates an otherwise less favourable outcome (epigenetic inheritance). Consistently, we observed that the levels of epigenetic silencing at *p53* was progressively enhanced during successive days *after* DOX removal (Fig. 5B). Moreover, this situation would suggest epigenetic inheritance in wildtype cells is highly genomic context-dependent. To test this, we targeted heterochromatin to multiple additional loci and tracked memory in endoderm. We observed that most regions reverted, albeit at least two (*Greb1* and *Jade1*) exhibited robust inheritance of a prior silenced state in wildtype endoderm (Fig S6C). Taken together, these data suggest that the potential for epiallele inheritance of *de novo* heterochromatin is influenced by multiple cell-type- and genomic-context-dependent factors. In the case of *p53*, augmenting weak-acting heterochromatin inheritance in differentiated with a favourable advantage changes the balance of dynamic forces to enable propagation of ectopic chromatin states within the population.

### Epigenetic inheritance during mammalian development in vivo

To more closely model the developmental process that occurs *in vivo* when pluripotent cells differentiate into all lineages, we differentiated ESC towards fates representative of multiple germ layers: ectoderm, endoderm, and epiblast like-cells (EpiLC) (as a model of primed pluripotency). Upon release of the iCRUSH heterochromatic trigger (Fig S7A), we observed maintenance of *p53* silencing in all three differentiation programs (Fig S7B). Notably this included primed EpiLC, emphasising the preferential capacity of naïve pluripotency to reset epialleles. All differentiating lineages replicated with comparable kinetics (Fig S7C). These data suggest that epigenetic aberrations acquired during the pluripotency window can potentially be inherited in all tissues.

**Figure 5.**
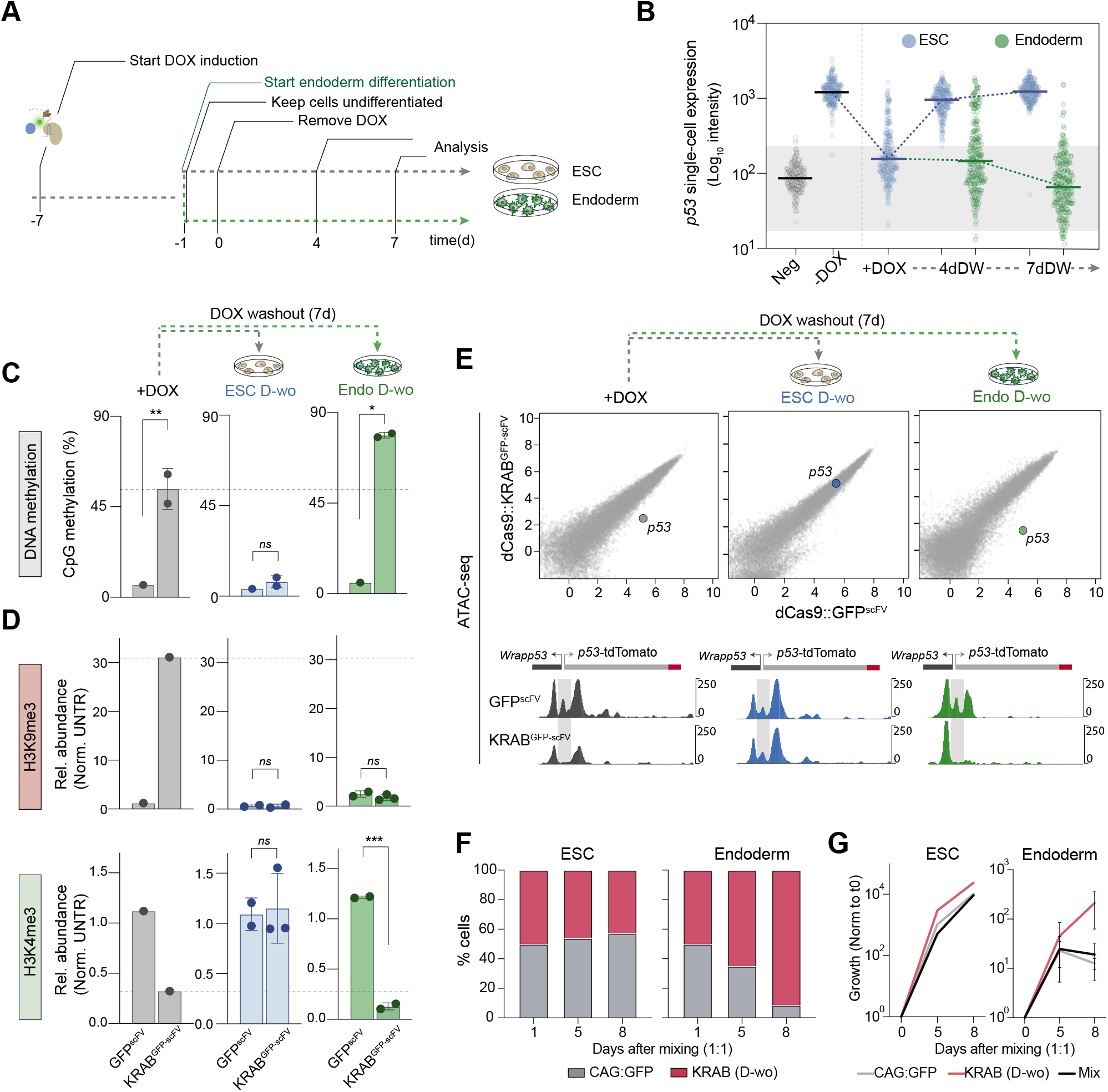
Aberrant epialleles can be propagated in wildtype cells upon exit from pluripotency. **(A**) Schematic of experimental timeline: heterochromatin epialleles are induced at *p53^-tdTomato^* in ESC followed by endoderm differentiation (day −1), with DOX removed after 24 hours (day 0). In parallel cells are maintained as naïve ESC. Chromatin and expression analysis to record memory in endoderm and ESC is performed at 4 and 7 days of DOX-washout. (**B**) Single-cell expression of *p53^-tdTomato^* during DOX-washout in endoderm or naïve ESC, following induction of heterochromatic silencing. (**C**) Bisulfite pyrosequencing quantification of DNA methylation at the *p53*-promoter in ESC or endoderm following DOX withdrawal. Dashed line indicates DNA methylation after targeting KRAB^GFP-scFv^ (+DOX) for 7d. (**D**) CUT&RUN-qPCR quantification of the relative memory of induced H3K9me3 and H3K4me3 at *p53* promoter in independent biological replicates of endoderm or ESC. (**E**) ATAC-seq scatterplots showing genome accessibility at all promoters comparing GFP^scFv^ and KRAB^GFP-scFv^ in +DOX or DOX washout conditions in ESC or endoderm cells. Shown below are relevant genome tracks of the *p53* promoter (highlighted in grey). (**F**) Ratio of KRAB^-GFP-scFv^ (D-wo) silenced cells (red) and CAG:GFP cells (grey) in ESC or endoderm after the indicated days from mixing 1:1. (**G**) Line plots indicate exponential growth of CAG:GFP, KRAB^-GFP-scFv^ (D-wo) or the mixed population in ESC or endoderm cells.

To determine whether perturbed epigenetic states acquired at loci such as *p53* can self-propagate during *in vivo* development, potentially affecting disease risk, we tested epigenetic inheritance during embryogenesis. We introduced *p53*-epigenetically silenced ESC (KRAB^GFP-scFv^) into E3.5 blastocysts and traced the memory during post-implantation development (-DOX), as compared to a control (GFP^scFv^) (Fig. 6A). By epifluorescence microscopy we observed a strong contribution of ESC to all tissues of the E10.5 embryo (Fig 6A), and a consistent fraction of the cells carried BFP expression analysed by flow cytometry (Fig 6C). Analysis of tdTomato within the BFP^pos^ cells, revealed *p53* is fully activated in controls, as expected (Fig 6C). In contrast, embryos with prior *p53* epigenetic silencing had a significant tendency (*p=0.03*) to propagate memory of this through development (Fig. 6D). Indeed, up to 7% of foetal cells inherited epigenetic silencing memory (Fig. 6D-E). Given the central role of *p53* as a tumour-suppressor, this has the potential to have a major impact on disease susceptibility.

Overall, these data suggest that an epiallele acquired during or after pluripotent phases can be inherited through subsequent development. Importantly, this effect is highly context-dependent and relies on supporting activities or influences that reinforce or promote propagation, either directly or indirectly.

**Figure 6.**
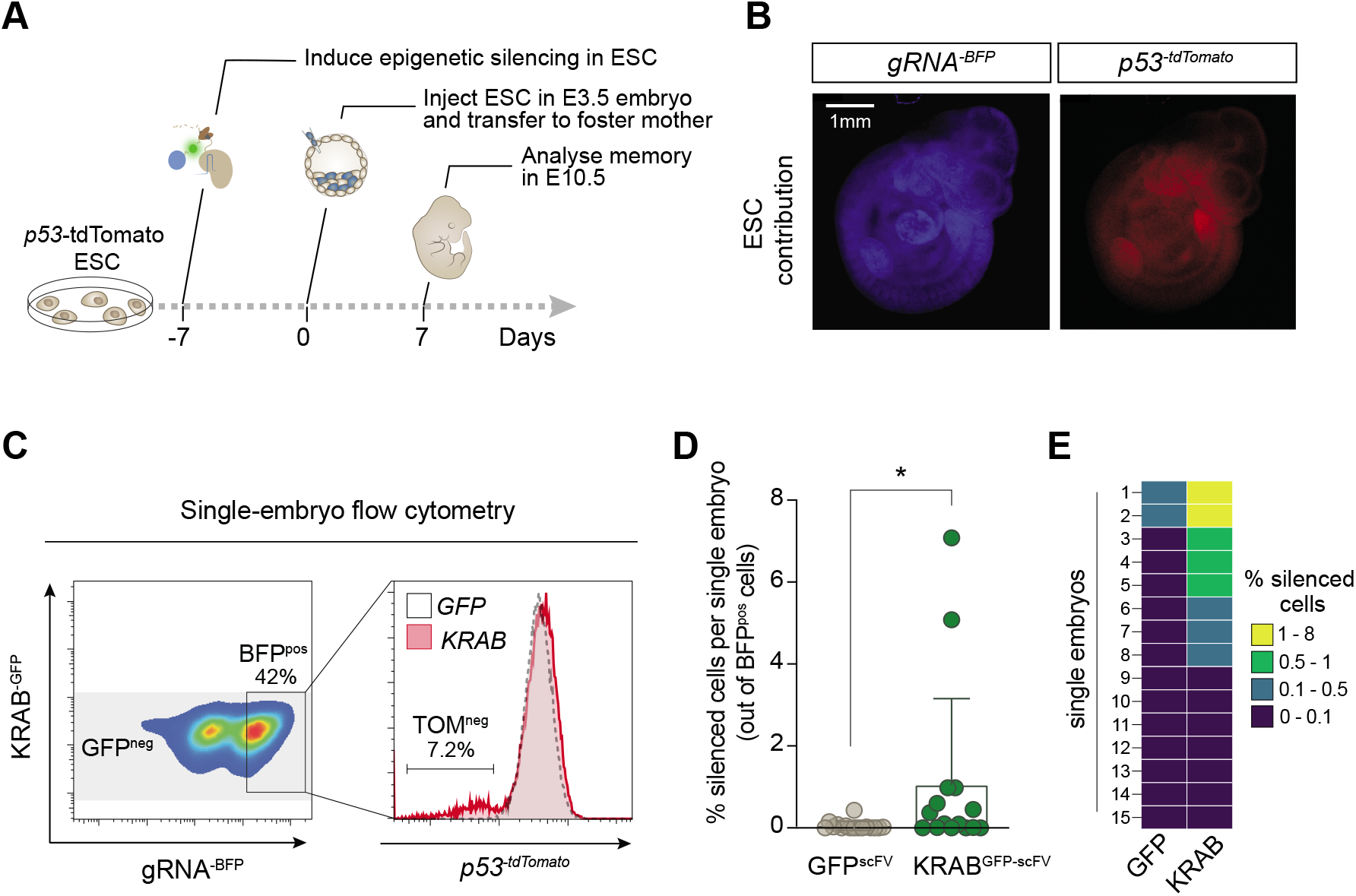
Induced epialleles exhibit epigenetic inheritance during *in vivo* development. (**A**) Schematic of experimental strategy to trace induced epialleles during in vivo development. (**B**) Epifluorescence images showing good contribution of ESC (marked in BFP expression) to E10.5 chimeric embryos. (**C**) Left: flow-cytometry density plot of gRNA^BFP^ and KRAB^GFP-scFv^ expression of a representative E10.5 embryo. Shown right is the distribution of *p53* expression in embryos demonstrating memory of induced *p53* silencing (KRAB) relative to control (GFP), in cells gated for BFP positive and GFP negative. (**D**) Percentage of *p53*-silenced cells in BFP^pos^/GFP^neg^ population. Each dot represents a single embryo from two independent experiments. (**E**) Heatmap showing the percentage of cells in each embryo that retained epigenetic silencing memory of *p53*.

## DISCUSSION

Here we used a precision epigenetic editing strategy (iCRUSH) to define the transcriptional function and memory of heterochromatin epialleles at endogenous loci. Our method of compound recruitment of multiple ‘effector’ modules using dCas9^GCN4^, facilitated programming of major (>10kb) heterochromatin domains, sufficient to drive robust epigenetic silencing. These *de novo* domains comprised H3K9me3, H4K20me3 and DNA methylation, and concomitant loss of H3K4me3, with modification levels comparable or greater than endogenous heterochromatic regions, which are thought to self-propagate via ‘read-write’ reinforcement (Reinberg & Vales, 2018).

Nevertheless, we found that naïve pluripotent cells act as a fundamental roadblock to inheritance of heterochromatin domains occurring outside of normal genomic contexts, even when providing a selective advantage such as silencing *p53*. This supports the concept of an epigenetic *‘tabula rasa’* during early mammalian development that acts to prevent intergenerational transmission of inherited or acquired chromatin epialleles. An exception to this principle is imprinted regions, wherein programmed chromatin was stably maintained, highlighting the role of the underlying DNA sequence context for epigenetic memory. This contextual influence is further exemplified by the effects of cell identity, with our data revealing the potential for epigenetic inheritance in mammals during normal *in vivo* development. Thus, we propose a unique and defining feature of naïve pluripotency is to reset aberrant chromatin states at endogenous loci to establish a pristine epigenome for development. Indeed, the functional properties of pluripotency *per se* are relatively unaffected by impairing global DNA demethylation (McLaughlin *et al,* 2019), and thus purging otherwise heritable epialleles could be a key operative function of epigenome reorganisation during pluripotent phases.

To decipher the underlying mechanisms that restrict epigenetic inheritance in naïve pluripotent cells, we designed a genome-wide CRISPR screening strategy. We found that loss of *Kmt2d* enables prolonged memory of epigenetic silencing in naïve ESC, presumably because of reduced H3K4me3 redeposition, yet the original epigenetic state is eventually restored, suggesting an important but non-critical role of *Kmt2d* in opposing heritable silencing. In contrast, we show that deletion of *Dppa2* enables robust long-term epigenetic inheritance of programmed epigenetic silencing in naïve cells. Interestingly this was a probabilistic effect, with most cells potentiating memory, but with a fraction delaminating to reactivate expression over time. This suggests that removing *Dppa2* shifts the balance of opposing factors to favor propagation of epigenetic silencing without fully saturating the odds against reversion.

Importantly, in *Dppa2* mutants the majority of loci remain in their erstwhile epigenetic configuration prior to acquiring a forced epimutation. This argues that loss of *Dppa2* sensitizes the genome to stably inherit any stochastic or programmed epigenetic changes, and implies DPPA2 acts as a general epigenome ‘surveyor’ to counteract epigenetic inheritance during pluripotent phases. This extends recent observations which showed a small subset of developmental loci and LINE1 directly acquire silencing in *Dppa2*-mutants (Gretarsson & Hackett, 2020), by revealing that loss of *Dppa2* renders a further fraction of the genome predisposed to inherit prospective epigenetic perturbations, potentially induced by external exposures. Mechanistically, DPPA2 is thought to target H3K4me3 through interactions with KMT2D (Eckersley-Maslin *et al.*, 2020), also a hit in our screen, which emphasizes that the molecular pathways that impair heterochromatin inheritance in ESC could converge on promoting antagonistic H3K4me3. Likely there are additional or redundant ‘surveyor’ factors since we observed epigenetic reversion of some loci in *Dppa2* mutants. This suggests there is a broad network of mechanisms that precludes epigenetic inheritance specifically in naïve pluripotent cells, with DPPA2, which is exclusively expressed during pluripotency, playing a central role.

In contrast to naïve cells, heritable epigenetic silencing has been reported in differentiated cells (Amabile *et al.*, 2016; Bintu *et al.*, 2016; Nunez *et al.*, 2021). This has however typically been in cancer-derived cell lines and the potential for mitotic propagation of *de novo* epigenetic states in a normal developmental context is relatively unexplored. We reasoned that if an epimutation occurred during or after early pluripotent phases *in vivo*, it could heritably influence subsequent cellular/organismal phenotype, via clonal inheritance in neighboring cells. Indeed, we found that whilst epigenetic silencing at some loci is reset, others demonstrate robust mitotic transmission of heterochromatin-mediated silencing, including *p53*. This appears to reflect the confluence of weak-acting probabilistic inheritance and a selective advantage conferred by stable *p53* repression, a phenomenon recently shown in yeast and dubbed Mendelian epigenetic inheritance (Catania *et al,* 2020; Torres-Garcia *et al.*, 2020). Importantly, we demonstrate this effect *in vivo* in mice, with up to 7% of cells within a whole embryo heritably maintaining the legacy of prior *p53* silencing. This is relevant as even a small fraction of organismal cells corresponds to a large absolute number that carry epigenetic silencing, and such constitutional epimutations have been linked to cancer risk (Hitchins, 2015). Moreover, the principle of probabilistic inheritance of epigenetic silencing *in vivo* shown here could have implications for other aspects of health and disease linked with early life environmental exposures that can promote epigenome changes.

In summary, we find the window of naïve pluripotency robustly counteracts induced epigenetic memory, in part through DPPA2 activity, implying an intrinsic role of naïve status is to erase epimutations during early mammalian development. Upon differentiation however, when *Dppa2* is downregulated, we find acquired chromatin states can self-propagate through development, particularly when providing a selective advantage. This highlights a previously unappreciated but crucial developmental function of naïve pluripotency, and paves the way for understanding the complex inputs that feed into epiallele propagation in mammals.

## ACKNOWLDEGMENTS

We are grateful to Marzia Munafò for critical reading of the manuscript and Monica Di Giacomo for experimental assistance. We are also grateful to all EMBL core facilities, in particular to Neil Humphries (GEEF) and Jim Sawitzke (GEVF), for key experimental support. We thank Claire Rougeulle, Sylvia Erhardt, Eileen Furlong and Mathieu Boulard for their contributions to thesis advisory committee discussions (for V.C). This study was funded by a European Molecular Biology Laboratory (EMBL) programme grant to J.A.H.

## AUTHOUR CONTRIBUTIONS

V.C performed experiments and analysis, and co-wrote the manuscript. C.P. provided experimental support. J.A.H designed and supervised the study, and wrote the manuscript.

## COMPETING INTERSTS

We declare no financial or non-financial competing interests.

## MATERIALS AND METHODS

### Routine cell culture

Naïve murine embryonic stem cells (mESC) were derived freshly (mixed 129/B6, XY), or obtained from (Hackett *et al.*, 2018), and were routinely cultured on gelatin-coated plates in t2i/L media: NDiff (N2B27) (Takara #y40002) supplemented with titrated 2i (0.2μM PD0325901, 3μM CHIR 99021), 1000 U/ml LIF, 1% FBS, 1% Pennicillin streptavidin and mainatained in humified atmosphere at 37°C and 5% CO2. Cells were passaged every 2-3 days by dissociation with TrypLE and medium was changed daily. Mycoplasma contamination checks were performed routinely by ultrasensitive qPCR assay (Eurofins).

### DNA transfection

DNA transfection was accomplished with Lipofectamine 3000 (Thermo Fisher Scietific #L3000001) unless otherwise stated. ES Cells were seeded at least 24 hrs in advance to be ~50% confluent the day of transfection. Appropriate amounts of DNA were calculated according to manufacturer’s instructions. Media was changed after 6 hours, and replaced with antibiotic selection containing medium where appropriate.

### Flow cytometry

For fluorescence activated cell sorting (FACS) or flow analysis, cells were gently dissociated in cell suspension by TrpLE, resuspended in PBS plus FBS 1% (FACS media) and filtered (BD, cup-Filcons #340632). A FACS Aria III (Becton Dickinson) or Attune NxT Flow Cytometer (Thermo Fisher Scientific) were used for sorting or analysis respectively. Data analysis was performed with FlowJo v10.5.3 (Tree Star, Inc.).

### Epigenetic editing tool constructs

Epigenetic editing tools comprising dCas9^GCN4^, KRAB^GFP-scFV^, 3a3L^GFP-scFv^ and GFP^scFV^ were cloned into PiggyBac recipient plasmids, properly linearized with restriction enzymes, by homology arm recombination using In-fusion HD-Cloning (Takara #639650) according to manufacturer’s instructions (Fig. S1A). For the pPB_TRE3G::dCas9^-5XGCN4^_*EF1a*::TetOn-Hygro, the *Streptococcus pyogenes* dCas9^GCN4^ was PCR amplified from the PlatTET-gRNA2 plasmid (Morita et al., 2016) (Addgene #82559), and cloned together with a d2 destabilization domain under control of the TRE3G promoter in a PiggyBac backbone vector also containing the TET-ON3G transactivator and the Hygromycin resistance gene separated by an IRES sequence and under control of the CAG promoter. For the effector plasmids (pPB_TRE3G::ScFv-GFP-KRAB_EF1a::Neo and pPB_TRE3G::ScFv-GFP-3a3L_EF1a::Neo) the GCN4 specific scFv domain and the sfGFP gene were amplified from PlatTET-gRNA2 plasmid (Addgene #82559) and fused in frame with the human ZNF10 KRAB domain (amplified from the pAAVS1-NDi-CRISPRi (Addgene #73498)) or the catalytic domain (CD) of mouse Dnmt3a and the C-terminal part of mouse Dnmt3L (3a3L) (amplified from pET28-Dnmt3a3L-sc27 (Addgene #71827)) and cloned in PiggyBac plasmids under control of the TRE3G promoter. The effector is also destabilized with a d2 domain and the vector also carries constitutive expression of the Neomycin resistance gene. The control effector GFP^scFV^ was cloned as described above but without any epigenetic domain. Finally, abolishment of the cytosine methyltransferase catalytic activity in Dnmt3a CD (Mut3a3L) was achieved by a single replacement of a cysteine at position 296 by a serine (Hsieh, 1999).

Similarly, the U6::gRNA_EF1a::BFP-Puro, carrying an enhacend gRNA scaffold, was amplified from Addgene plasmid #60955 and cloned into into a PiggyBac recipient plasmid (pPB_U6::gRNA_EF1a::BFP-Puro).

To design all sgRNA for targeting the epigenetic editing system, the GPP web portal (Broad Institute) was used. Reverse complement gRNA sequences (Table 1) with appropriate overhangs were annealed at 10 μM final concentration with 10 mM Tris, pH 7.5 – 8.0, 60 mM NaCl, 1 mM EDTA, by heating at 95°C for 3 minutes and cooling down at RT for >30 minutes. Annealed sgRNA were ligated with T4-DNA ligase (NEB #M0202S) for 1 hour at 37°C into pPB_U6::gRNA_EF1a::BFP-Puro digested with BlpI (NEB #R0585S) and BstXI (NEB #R0113S). All ligated assembled plasmids were amplified by bacteria transformation and purified by endotoxin-free midi-preparations (ZymoResearch # D4200). Correct assembly and sequences were confirmed by Sanger sequencing (Genewiz).

### Generation of reporter cell lines

The *Esg1^-tdTomato^* reporter cell line was derived from the *Stella*-<u>G</u>FP:*<u>Esg1</u>*-tdTomato (SGET) compound-reporter line (Hackett *et al.*, 2018). For generating the *p53^-tdTomato^* reporter cell line, we obtained T2A-tdTomato dsDNA sequence by PCR amplification from a donor vector with ultramers carrying 180 bp overhangs complementary to the 3’ end of the *p53* gene. We introduced this into cells by transfection of 129/B6 XY ESC together with the spCas9 plasmid pX459 (Addgene #62988), carrying a single gRNA sequence complementary for the *p53* 3’-end. After antibiotic selection for transient px459, TOM^pos^ single cells were sorted at by FACS. Single cells were expanded clonally and correct mono-allelic integration to generate *p53^-tdTomato^* was validated by PCR genotyping and Sanger sequencing (Genewiz). Normal levels of *p53* mRNA expression were verified by qPCR.

### Epigenetic editing and memory assay

For stable integration of the epigenetic editing system *Esg1^-tdTomato^* or *p53^-tdTomato^* WT or KO reporter ESC lines were co-transfected with the Piggybac plasmids: pPB_TRE3G::dCas9^-5XGCN4^_*EF1a*::TetOn-Hygro, pPB_TRE3G::ScFv-KRAB^-GFP^_EF1a::Neo, pPB_U6::gRNA_EF1a::BFP-Puro containing appropriate gRNA sequence and pPY_CAG_Pbase using 5:5:1:1 molar ratio, respectively. Alternatively the pPB_TRE3G::KRAB^-GFP-scFv^_EF1a::Neo was replaced with a construct carrying only expression of *GFP^-scFv^* pPB_TRE3G::ScFv-GFP_EF1a::Neo as a control. Cells with successful integration of the 3 cassettes were enriched by successive selection with hygromycin (250 μg /ml) for 5 days, neomycin (300 μg/ml) for 3 days and puromycin (1.2 μg /ml) for 2 days. After two days of cellular recovery, expression of dCas9^5XGCN4^ and *KRAB^-GFP-scFv^* was induced with Doxycyclin (DOX) (100 ng/ml) for 7 days, and double GFP and BFP positive cells, that had activated the epigenetic editing system, were enriched by FACS. GFP/BFP double positive cells were re-seeded into culture in absence of DOX and retention of reporter silencing estimated by flow cytometry after 4 or 7 days of DOX washout in cells that switched off the destabilised epigenetic editing tool gated as BF^pos^/GFP^neg^ cells.

### Cell cycle inhibition

For the cell cycle inhibition to test active vs passive epigenetic erasure, the *Esg1^-tdTomato^* cell line, already carrying the *dCas9^-5XGCN4^* and *KRAB^-GFP-scFv^* or *GFP^-scFv^* were transiently transfected with a pPB_U6::gRNA_EF1a::BFP-Puro containing gRNA against *Esg1* TSS (*gRNA^87dw^*). 1.2 ng/ml of puromycin selection was added after 6 hours together with DOX (100 ng/ml). Cells were cultured for 3 days and then sorted for TOM^neg^ status. TOM^neg^ cells were then cultured in absence of DOX for a total of four days, with the cell cycle inhibitor RO3306 (9 μM) added after 24 hours (when GFP had just switched off), and removed after 48 hours. Cells were analysed by flow cytometry at 24 hours intervals and gated for absence of expression of GFP and BFP.

### ESC differentiation

To induce endodermal, ectodermal or epiblast-like cell (EpiLC) differentiation, naïve ESCs were cultured for five days in presence of DOX (100 ng/ml) seeded at a confluence of 6×10^3^/cm^2^ on gelatin-coated plates (for endoderm and ectoderm) or fibronectin-coated plates (for EpiLC) and maintained in t2i/L media for 24 hours in presence of Doxycycline (100 ng/ml). After removal of ESC medium and 5x washes with PBS, differentiation was induced as follows: (i) endoderm differentiation was induced with IDE1 (STEMCELL Technologies #72512) -containing medium (Borowiak et al., 2009) (RPMI (Thermo Fisher Scietific #12-633-012) supplemented with 0.02% FBS, 2mM L-Glutamine, 5 μM IDE1 and 1% Pennicillin streptavidin); (ii) ectoderm differentiation was induced with NDiff (NB27 Takara #y40002) supplemented with 0,25 μM Retinoic Acid (Sigma Aldrich#R2625), 0,02% FBS and 1% Pennicillin streptavidin; (iii) EpiLC differentiation was induced with NDiff supplemented with 20 ng/ml ActivinA (PeproTech #120-14P), 12ng/ml bFGF (PeproTech #450-33), 1% knockout serum replacement (Thermo Fisher #10828010), and 1% Pennicillin streptavidin. In all cases, DOX treatment was maintained for the first 24 hours of differentiation and then cells washed 5 times and cultured for a maximum of 8 days without DOX, with media change every other day. Flow cytometry was performed at 5 and 8 days of differentiation (corresponding to 4 and 7 days of DOX washout respectively) and, at the same time, cells harvested for bisulfite pyrosequencing and CUT&RUN.

### Growth competition assay

For assaying the competition advantage of the *p53* epigenetically silenced cells, *p53^-tdTomato^* reporter line, carrying *dCas9^-5XGCN4^, KRAB^-GFP-scFv^* and a gRNA against p53 (*gRNA^345up^*), were induced with DOX (100 ng/ml). In parallel, *p53^-tdTomato^* reporter line without the epigenetic editing tool from a similar passage number, were transfected with a PiggyBac plasmid driving constitutive GFP expression (CAG:GFP) and subjected to two subsequent rounds of sorting to enrich GFP^pos^ cells. After 7 days of DOX induction, KRAB-induced *p53^-tdTOM^* negative (TOM^neg^) cells were enriched by flow cytometry and equally mixed (1:1) with cells expressing constitutive GFP and subjected to DOX washout. After 5 or 8 days from mixing, cells were analysed by flow cytometry to measure the proportion of GFP^pos^ and GFP^neg^ cells.

### Generation of knockout ESC lines

Knockouts (KO) cell lines were generated by transiently transfect two spCas9 plasmids (pX459) carrying one gRNA each targeting exon sequence for critical catalytic activity of the gene of interest (*Dppa2, Zmym3, Kmt2d, Smarcc1, Dot1L*) (Table 1) in *Esg1^-tdTomato^* reporter lines. After transfection, cells were selected with puromycin (1.2 ug/ml) for three days and subsequently seeded at low density (1000 cells per 9.6cm^2^) for single colony picking. Subsequently to expansion, single clones were screened for bi-allelic genetic deletion by PCR genotyping (Table 2) and sanger sequencing (Genewiz). For *Dppa2^-/-^*, absence of the protein was further confirmed by western blot.

### Western Blot

For protein extraction, cell pellets were resuspended in RIPA buffer (Sigma #R0278) containing protease inhibitors (Roche #4693159001), incubated at 4°C for 30 minutes and, upon centrifugation, cell lysis supernatant was collected. 10-20 μg of proteins were mixed with bolt LDS sample buffer (ThermoFisher #B0007) and bolt reducing agent (ThermoFisher #B0004), heated at 70°C for 10 minutes and loaded on 4-12% Bis-Tris gel (ThermoFisher #NW04125BOX). After electrophoresis separation at 200V using MES running buffer (ThermoFisher #NP0002), proteins were transferred to a PVDF membrane (ThermoFisher #IB24002) using the iBlot 2 transfer stack (ThermoFisher) and the membrane was subsequently saturated with with 5% milk/1xPBS for one hour at room temperature. For detection of the protein of interest, the membrane was incubated at 4°C overnight with primary antibody (Table 3) in 0.5%milk/PBS/0.05%tween and after three washes in PBS/0.05% tween, HRP-linked secondary antibody incubation was carried on in 0.5%milk/PBS/0.05%tween for 1 hour at room temperature. After washing thrice with 0.5%milk/PBS/0.05%tween, detection was performed by incubating the membrane with Pierce ECL western blot solution (ThermoFisher #32132) for 5 minutes prior imaging with ChemiDoc XRS+ system (BioRad).

### RNA preparation and real-time qPCR

Total RNA was extracted using the PicoPure RNA isolation kit (Applied Biosystems #KIT0204) for less than 1×10^4^ cells or the RNeasy kit (Qiagen #74004) otherwise, following manufacturer instructions. 1μg of RNA was used as input to generate complementary DNA (cDNA), with a mixture of random hexamers and reverse transcriptase, following DNAase treatment (TAKARA PrimeScript RT Reagent Kit with gDNA Eraser #RR047A). A control reaction in which the RNA was incubated with all the other components except the reverse transcriptase enzyme mix (-RT control) was performed. cDNA was diluted 1:10 and specific targets quantified by real-time quantitative qPCR using primers designed at exon-exon junctions to minimise amplification from contaminant DNA (Table 2). The reaction was performed using SYgreen Blue Mix (PCRbio # PB20.15-20) and a QuantStudio 5 (Applied Biosystems) thermal cycler.

### Bisulfite pyrosequencing

DNA bisulfite conversion was performed directly starting from cell pellets (a maximum of 1×10^5^ cells per sample) using the EZ DNA Methylation-Direct kit (Zymo Research #D5021) following the manufacturer’s instructions. Target genomic regions were PCR amplified using 1μl of converted DNA with biotin-conjugated bisulfite primers (Table 3), using the PyroMark PCR kit (Qiagen #978703). Pyrosequencing assay conditions were generated using the PyroMark Q24 Advanced 3.0 software and the sequencing reaction was performed with PyroMark Q24 advanced reagents (Qiagen, #970902) according to manufacturer’s instructions. Briefly, 10 μl of the PCR reaction was mixed with streptavidin beads (GE Healthcare #17-5113-01) by shaking for 5 minutes at room temperature and, after separation of DNA strands and release of samples into the Q24 plate (Qiagen) using PyroMark workstation (Qiagen), sequencing primers were annealed to DNA by heating at 80°C for 2 minutes and cooling down at RT for 5 minutes. Pyrosequencing was run on PyroMark Q24 advanced pyrosequencer (Qiagen) with target specific dispensation order (Table 4). Results were analysed with PyroMark Q24 Advanced 3.0 software.

### CUT&RUN

The CUT&RUN (Cleavage Under Targets and Release Using Nuclease) protocol (Skene & Henikoff, 2017) was used to detect protein-DNA interaction and histone modifications. Briefly, a total of 3×10^5^ cells per sample were pelleted and washed twice with wash buffer (20mM HEPES pH 7.5, 150mM NaCl, 0.5 mM Spermidine containing protease inhibitor) and incubated with conacavallin A magnetic beads (Sigma Aldrich C7555) by rotating for 10 minutes at room temperature. After placing samples on a magnet stand, the supernatant was removed. Cells were resuspended with antibody buffer (wash buffer with 0.02% digitonin, 2mM EDTA) containing 0.5 μg of target-specific antibody (Table 3), and left rotating overnight at 4°C.

Samples were then placed on a magnet stand to remove antibody buffer, washed thrice with wash buffer containing 0.02% digitonin (Dig-wash buffer), and incubated with 700 ng/ml of purified protein-A::MNase fusion (pA-MNase) on a rotor at 4°C for one hour followed by two more washes. MNase reaction was thus activated by adding 4mM CaCl_2_ and incubating at 0°C for 30 minutes and immediately stopped with 1X final concentration of STOP buffer (340mM NaCl, 20mM EDTA, 200mM EGTA, 0.02% Digitonin, 250 μg glycogen, and 250 μg RNaseA). Target chromatin was released by incubating at 37°C for 10 minutes, centrifuging at full speed for 5 minutes at 4°C and the supernatant collected after incubation on magnet stand. DNA was finally released from chromatin by incubation with 0.4% SDS (Promega #V6551) and 0,5 mg/ml Proteinase K (Thermo Fisher Scientific #AM2546) at 70°C for 10 minutes. Purification and size selection of DNA was performed using SPRI beads (Beckman Coulter #B23318) following the instruction for double size selection with 0.5X and 1.3X bead volume to sample volume ratio. CUT&RUN DNA fragments were either subjected to quantitative-PCR to amplify selected targets or to next generation sequencing to quantify chromatin/marks abundances genome/wide.

For CUT&RUN-qPCR, DNA fragments were diluted ten times with H_2_O and 2 μl amplified with SYgreen Blue Mix (PCRbio) and primers specific for target and control regions (in which the mark is expected to be enriched (positive controls) or depleted (negative controls)) (Table 2) using the QuantStudio 5 (Applied Biosystems) thermal cycler. Note that primers were designed to amplify minimum amplicon sizes as CUT&RUN produces small fragments. Relative abundance of histone marks was estimated comparing Ct-values of target regions to positive control regions.

For CUT&RUN-sequencing libraries were made starting from 10 ng of CUT&RUN DNA fragments using the NEBNext Ultra II DNA Library Prep Kit for Illumina (NEB #E7645S) using the following PCR program: 98°C 30s, 98°C 10s, 65°C 10s and 65°C 5min, steps 2 and 3 repeated for 12-14 cycles depending on input DNA. After quantification and quality check with an automated electrophoresis system (Agilent Tape Station system), library samples were sequenced on the Nextseq Illumina sequencing system (paired-end 40 sequencing). Raw Fastq-sequences were trimmed to remove adaptors with TrimGalore (v0.4.3.1, -phred33 --quality 20 --stringency 1 -e 0.1 --length 20), quality checked and aligned to the custom mouse mm10 genome with the inserted tdTomato reporter using Bowtie2 (v2.3.4.2, -I 50 -X 800 --fr -N 0 -L 22 -i ‘S,1,1.15’ --n-ceil ‘L,0,0.15’ --dpad 15 --gbar 4 --end-to-end --score-min ‘L,-0.6,-0.6’). Analysis of the mapped sequences was performed using seqmonk (Babraham bioinformatics, v1.46.0) by enrichment quantification of the normalised reads.

### ATAC-seq

Prior to harvesting, cells were initially treated in culture medium with 200 U/ml of DNase for 30 minutes at 37°C to digest degraded DNA released from dead cells. After 5x washes with 1xPBS, cells were detached with TryPLE, 5×10^4^, counted and pelleted at 500 RCF at 4°C for 5 minutes. Supernatant was removed and cells resuspended in 50 μl of cold ATAC-Resuspension Buffer (10 mM Tris-HCl pH7.4, 10 mM NaCl, 3 mM MgCl2) with 0.1% NP40, 0.1% Tween20 and 0.01% Digitonin and incubated on ice for 3 minutes. Lysis was washed out using 1 ml of cold ATAC-Resuspension Buffer with 0.1% Tween20 and mixed. Nuclei were pelleted at 500 RCF for 10 minutes at 4°C. After removal of supernatant, nuclei were resuspended in 50 μl of transposition mixture (25 μl 2xTD buffer, 2.5 μl transposase (Illumina Tagment DNA Enzyme and Buffer Kit #20034197), 16.5 μl PBS1x, 0.5 μl 1% digitonin, 0.5 μl 10% tween20, 5 μl H_2_O) and incubated at 37°C for 30 minutes in a thermomixer with 1000 RPM shaking. Reaction product was cleaned-up with DNA clean and concentration kit (Zymo Research #D4014) following manufacturer instructions and eluted in 21 μl of elution buffer. 20 μl of this product was used for PCR amplification using Q5 Hot Start High-Fidelity polymerase (NEB #M0494S) and a unique combination of the dual barcoded primers P5 and P7 Nextera XT Index kit (Illumina #15055293) following the cycling conditions: 1. 98°C for 30 sec; 2.98°C for 10 sec; 3. 63°C for 30 sec; 4. 72°C for 1 min; 5. 72°C for 5 min and repeat from 2-4 for 5 cycles. After the first 5 cycles, 5 μl of the pre-amplified mixture was used to determine additional cycles by qPCR amplification using SYgreen Blue Mix (PCRbio) and the above used P5 and P7 primers in a QuantStudio 5 (Applied Biosystems) thermal cycler. After qPCR amplification, profiles were manually assessed plotting linear Rn versus cycle and the number of the additional PCR cycles to be performed equals to one-third of the maximum fluorescent intensity in this plot (Buenrostro *et al,* 2015). The identified number of extra PCR cycles were performed by placing the pre-amplified reaction back in the thermal cycler. Final clean-up of the amplified library was performed using the DNA clean and concentration kit (Zymo #D4014) and DNA amplicons eluted in 20 μl of H_2_O. After quantification and quality check with an automated electrophoresis system (Agilent Tape Station system), library samples were pooled together and sequenced on the Nextseq Illumina sequencing system (paired-end 40 sequencing).

For sequencing, raw reads were first trimmed with with TrimGalore (v0.4.3.1, reads>20bp, quality >30) and then quality checked with FastQC (v0.72). Output files were aligned to custom mouse mm10 genome with the inserted tdTomato reporter using Bowtie2 (v2.3.4.3, Paired-end settings, fragment size 0-1000, --fr, allow mate dovetailing). Uninformative reads were removed with Filter BAM (v2.4.1, mapQuality>=30, isProperPair, !chrM) and duplicated reads were filtered with MarkDuplicates tool (v2.18.2.2). The mapped and filtered sequences were then analysed with seqmonk (Babraham bioinformatics, v1.46.0) by performing enrichment quantification of the normalized reads. Correlation plots were generated by comparing enrichment of reads at promoters in sample versus control conditions.

### Genome-wide CRISPR screen and analysis

For the genome-wide screen, stable integration of *spCas9*-T2A-GFP was achieved in *Esg1^-tdTomato^* reporter ESC by insertion into the *Rosa26* safe harbour locus by CRISPR targeting with a pair of *Rosa26* specific gRNAs. After antibiotic selection and single-cell GFP^pos^ FACS sorting, integrity of the construct was verified by PCR genotyping and sanger sequencing. The PiggyBac *dCas9^GCN4^* construct was subsequently introduced in these cells as described before and after single colony picking and expansion, successful integration of *dCas9^-GCN4^* was functionally tested.

To introduce the genome wide perturbation, we used Lentiviral vectors carrying the Brie gRNA library (Doench et al., 2016) produced as previously described (Carlini *et al,* 2020). Briefly, the pooled gRNA Brie library (Addgene #73632) was expanded in HEK 293T according to BSL2 guidelines, lentiviral-containing supernatant was harvested and viral particles concentrated and resuspended in NDIFF 227. Lentiviral activity was estimated by transducing ESC across a titration curve and identifying a titration ratio to obtain 30-50% infection efficiency. 7×10^7^ *Esg1^-tdTomato^* ESC containing the CAG::spCas9-T2A-GFP and *TRE3G::dCas9^-GCN4^* were transduced in t2i/L medium with the pre-determined volume of lentiviral particles to ensure ~50% efficiency (>400 fold gRNA coverage). After 24 hours, we removed any residual lentiviral particles by five washes with PBS1X and we selected the cells using puromycin (1,2 μg/ml) for 7 days. Cells were then passaged before confluence, maintaining a minimum of 3.2×10^7^ cells to ensure gRNA library coverage (>400 fold coverage) and medium was changed daily for one week to give enough time for the knockouts to be generated.

Before introducing the epigenome editing tool we inactivated the spCas9-T2A-GFP cassette by transfecting >1×10^8^ KO pool cells with two tracr:crRNA (eurofins, Table 1) against two unique sequences in the *spCas9* that differs from *dCas9^GCN4^* using Xfect RNA transfection reagent (Takara #631450) according to manufacturer instructions. After five days from transfection, 3×10^7^ GFP^neg^ cells were sorted with FACS to select the cells with inactivation of the *spCas9*, plated back in t2i/L 10%FBS and further expanded for three days. To introduce the epigenetic perturbation, 2×10^8^ *Esg1^tdTomato^* KO library cell line already carrying dCas9^*GCN4*^ were transfected with *pPB_TRE3G::KRAB^-GFP-scFv^_EF1a::Neo, pPB_U6::gRNA_EF1a::BFP-Puro* containing a gRNA against *Esg1* and *pPY_CAG_Pbase* using Xfect mESC transfection reagent (Takara #631320) and selection (Neomycin (300 μg/ml) and DOX (100 ng/ml) induction was started after 24 hours. Seven days post-transfection, 3×10^7^ cells were sorted for TOM^neg^ and plated back in culture in absence of DOX. After 4 days of DOX washout, 3×10^7^ cells were sorted in parallel from the TOM^neg^ and TOM^pos^ fractions for genomic DNA extraction as an early timepoint (D-wo (3d)), using a gating strategy to separate fully silenced cells (TOM^neg-2.5%^) or cells ranging from fully to mildly silenced (TOM^neg-wide^). At the same time, 3×10^7^ unsorted cells were passaged up to a total of 7 days of DOX washout and sorting have been repeated as before to separate TOM ^neg-2.5%^ TOM^neg-wide^ and TOM^pos^ for the final timepoint (D-wo (7d)). Genomic DNA was isolated from purified populations by using DNeasy blood and tissue kit (Qiagen # 69504) following manufacturer instruction including RNAse step.

DNA libraries were prepared from TOM ^neg-2.5%^, TOM^neg-wide^ and TOM^pos^ at D-wo (3d) and D-wo (7d) timepoints in multiple parallel reactions each containing 500 ng of gDNA, with custom primers containing the P7 flow cell overhangs (5’-CAAGCAGAAGACGGCATACGAGATNNNNNNNNGTGACTGGAGTTCAGACGTGTGCTCTTCCGATCTTCTACTATTCTTTCCCCTGCACTGT-3’) including 8 bp barcode and P5 overhang (5’-AATGATACGGCGACCACCGAGATCTACACTCTTTCCCTACACGACGCTCTTCCATCTTTGTGGAAAGGACGAAA CACCG-3’) using the Q5 Hot Start High-Fidelity polymerase (NEB #M0494S) for 22-24 cycles. sgRNA amplicons were purified using SPRI beads (Beckman Coulter #B23318) following the instruction for double size selection with 0.5X and 1.2X bead volume to sample volume ratio. Purified fragments were checked and quantified with a tape station automated electrophoresis system (Agilent). Equal amplified library amounts were pooled together into a multiplexed library and sequenced for singleend 50bp (SE-50).

Counting of sgRNA representation in the isolated subpopulation of cells was performed using the Model-based Analysis of Genome-wide CRISPR-Cas9 Knockout (MAGeCK, v0.5.9) tool (Li, 2014). Briefly, the reads were first trimmed using cutadapt (v1.15) (*cutadapt -g TTGTGGAAAGGACGAAACACCG*) and quality checked using FastQC and then, the gRNAs counts were normalised to total reads within the sample (MAGeCK -count -norm-method total). Last, the TOM ^neg-2.5%^ or TOM^neg-wide^ were compared to the TOM^pos^ for each timepoint to identify significantly enriched/depleted gRNAs with a false discovery rate (FDR) < 0.2, using the -test command in MAGeCK.

### Embryo manipulation

Prior to microinjection, *p53^-tdTomato^* reporter ESC were transfected with dCas9^GCN4^, *p53_ gRNA^345up^*, and *KRAB^GFP-scFv^* or alternatively *GFP^-scFv^* and treated with DOX for 7 days. ESC microinjection was performed by the Gene Editing & Embryology Facility of EMBL Rome, using E3.5 embryos derived from natural mating of C57BL/6J mice. Injected embryos were implanted back into pseudo-pregnant foster mothers. All animals employed and procedures were in accordance with the gold-standard Italian and European Union regulation guidelines and approved by the local ethical committee.

After 7 days from the injection, at the embryonic development day 10.5, the deciduum was collected from the uterus and put into a 6 cm dish with cold PBS+10% FBS. Embryos were then extracted from the deciduum and moved to fresh PBS+10% FBS, placenta removed and cleaned from debrids and tissue fragments. Individual embryos were moved in one well of a round-bottom 96 well plates (Corning #CLS3367) containing 50μl of TripLE and incubated at 37°C for 20 minutes and pipetted until the embryo is entirely dissociated in single cell. The single cell suspension was then diluted with 100μL of PBS+1%FBS and spun down at 1200rpm for 5 min. Cell pellet was then resuspended in 300μl of FACS medium (1xPBS+1%FBS) and filtered (BD, cup-Filcons #340632) for quantitative flow cytometry analysis with Attune Nxt.

