## Supplementary Figures for "Epigenetic Inheritance is Gated by Naïve Pluripotency and *Dppa2*"

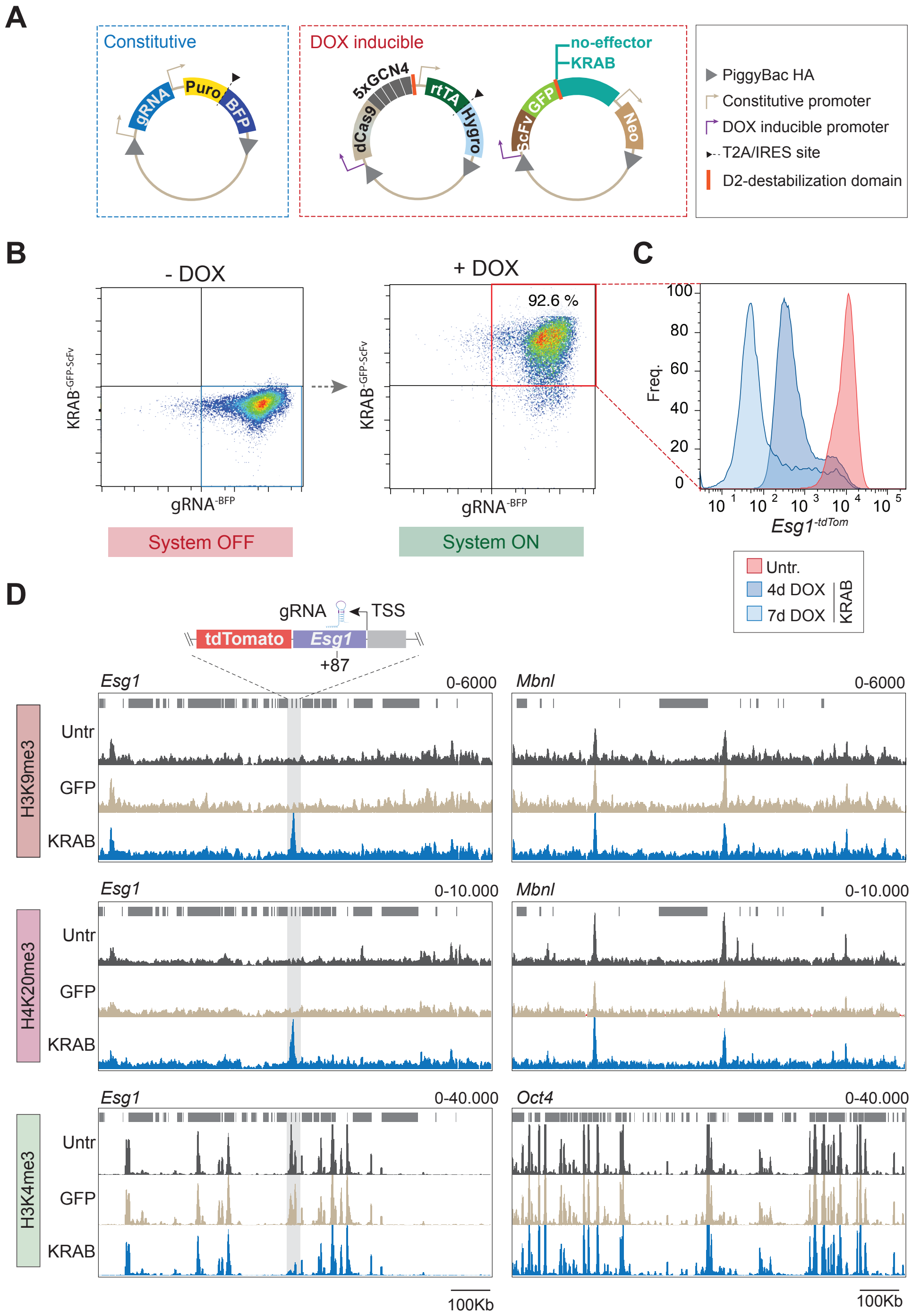


**Fig. S1 iCRUSH induces domains of *de novo* histone modification levels comparable to endogenous heterochromatin loci**

(**A**) PiggyBac constructs used to deliver the epigenetic editing tool (iCRUSH) into ESC. From left to right: the enhanced gRNA scaffold is under control of a constitutive U6 promoter, the same construct also drives constitutive expression of a puromycin selection gene and Blue Fluorescent Protein (BFP) separated by the self-cleaving peptide T2A; the *dCas9*$^{GCN4}$ gene is under control of the TRE3G DOX inducible promoter, the same construct also drives constitutive expression of the rtTA trans-activator and a hygromycin resistance gene separated by a T2A; the epigenetic effector construct (KRAB$^{GFP\text{-}scFV}$ or Dnmt3a3L $^{GFP\text{-}scFV}$) is also under control of the TRE3G DOX inducible promoter and it is fused with a Green Fluorescent Protein (GFP) and an scFv domain specific for GCN4, it also drives constitutive expression of the neomycin resistace gene. These constructs are genomically integrated into ESC by co-transfection with the piggybac transposase. (**B**) Density plots, obtained by flow cytometry analysis, show gRNA$^{BFP}$ and KRAB$^{GFP\text{-}scFv}$ expression in the cell population prior to and upon DOX induction, revealing its dynamic activation. (**C**) Fluorescence distribution measured by flow cytometry showing *Esg1*-reporter silencing after recruitment of KRAB$^{\text{-}GFP\text{-}scFv}$ for 4 or 7 days (blue) compared to a reference untargeted sample (red), demonstrating order of magnitude silencing. Cells were gated for the presence of both KRAB$^{scFv\text{-}GFP}$ (GFP positive) and gRNA$^{\text{-}BFP}$ (BFP positive). (**D**) Genome views comparing the magnitude of *de-novo* peaks of histone marks deposited by iCRUSH at the *Esg1* promoter (left) with endogenous representative examples (right).

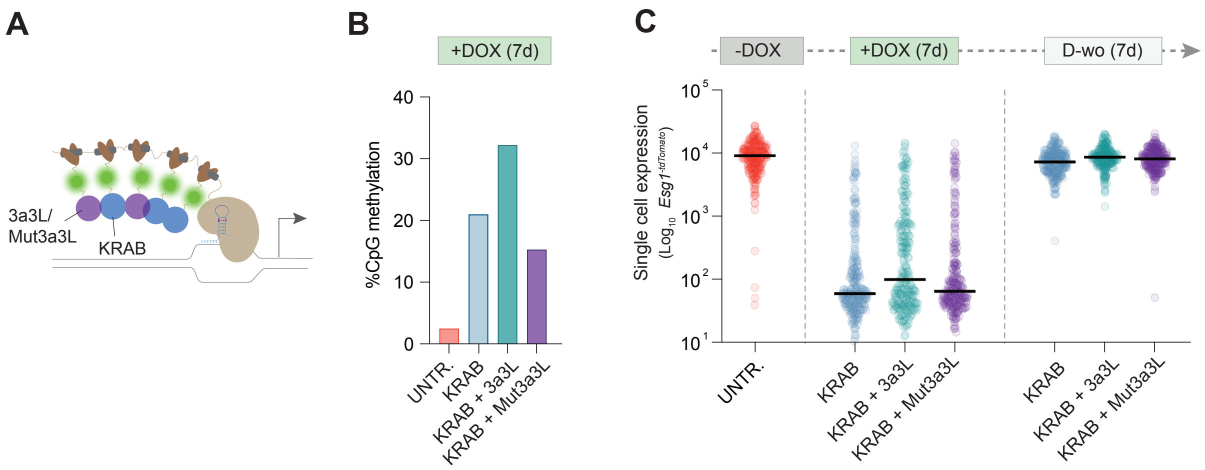


**Fig. S2. Combination of KRAB and Dnmt3a3L does not enhance epigenetic memory in naïve ESC**

(**A**) Schematic of the iCRUSH epigenetic tool used in the experiment. $KRAB^{GFP\text{-}scFv}$ was either recruited alone or in combination with Dnmt3a/Dnmt3L ($3a3L^{GFP\text{-}scFv}$) or the catalytically mutant $Mut3a3L^{GFP\text{-}scFV}$. (**B**) Histogram plot showing the average of percentage DNA methylation at *Esg1* promoter assayed by bisulfite pyrosequencing. (**C**) Violin plots show the single-cell distribution of $Esg1^{\text{-}tdTomato}$ expression (log scale) in each condition (-DOX, 7days +DOX and 7days DOX washout) upon epigenetic editing with $KRAB^{\text{-}GFP\text{-}scFv}$ (blue) +/- $3a3L^{GFP\text{-}scFv}$ (aqua green) or $Mut3a3L^{GFP\text{-}scFV}$ (purple).

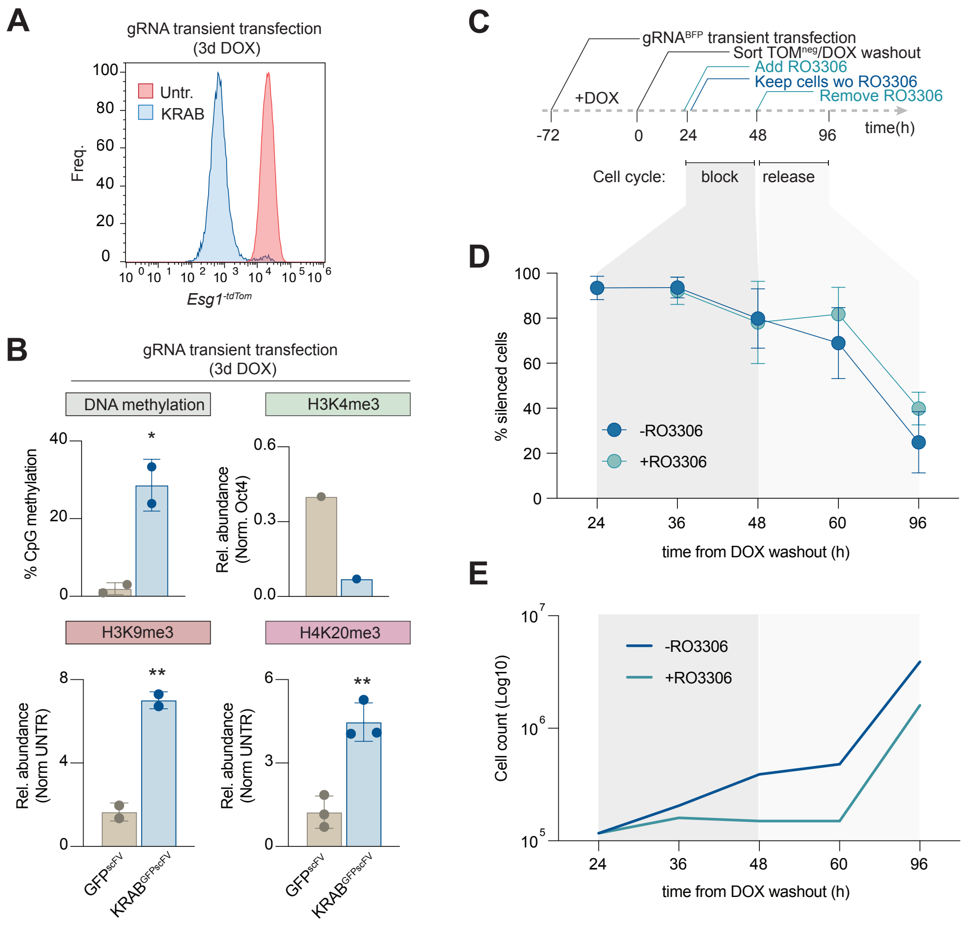


**Fig. S3. Cell cycle inhibition slows but does not block epigenetic erasure in naïve ESC.**
(**A**) *Esg1*$^{-tdTomato}$ reporter expression in KRAB$^{GFP\text{-}scFv}$ or untransfected control after gRNA transient transfection and 3 days of DOX induction. (**B**) Histograms show percentage of CpG methylation or relative abundance of histone modifications nomalised to a positive control region and untransfected control after 3 days of DOX (gRNA transiently transfected). Statistics calculated by one-tail unpaired t-test over two or three independent biological replicates *(*=p<0.05; **=p<0.01)*. (**C**) Timeline of the experiment. Briefly, cells are treated for 72 hours (3days) with DOX to induce iCRUSH mediated silencing. tdTomato negative cells are sorted and plated back in culture in duplicate expeirments without DOX. After 24 hours of DOX washout one sample is treated with the cell cycle inhibitor RO3306 for further 24 hours. After a total of 48 hours from DOX washout the cell cycle is released and cells further analysed at 12 hours interval up to 96 hours. In parallel cells are cultured in the absence of the inhibitor. (**D**) Time-course of the percentage of *Esg1*$^{tdTomato}$ silenced cells in the + or -RO3306 conditions at 12 hours intervals of DOX washout. Error is measured as standard deviation between two biological replicates. (**E**) Line plots indicate log growth of cells in + or – RO3306 conditions.

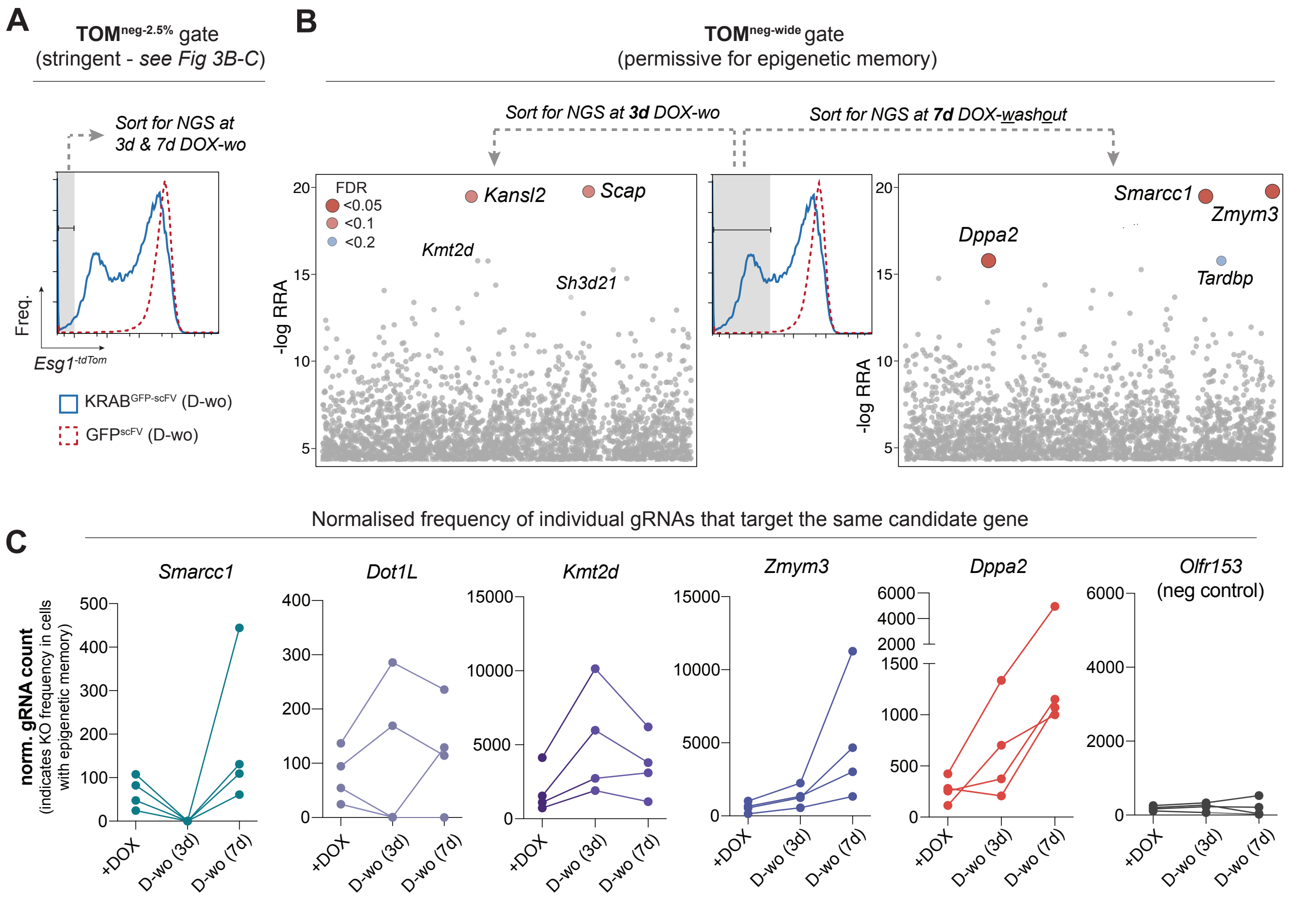


**Fig. S4. Gating strategy for CRISPR screen and supplementary scatterplots of candidate factors that antagonise epigenetic inheritance.** (**A**) Histogram showing the gate used to sort cells that strictly retained epigenetic silencing memory in the CRISPR screen (tdTomato-negative) using a stringent gating threshold (bottom 2.5% TOM- cells (TOM$^{2.5\%neg}$). The results of significant candidate genes that enable this memory when knocked-out are detailed in main Fig 3B-C. (**B**) Middle histogram: A wider gate was used to include all tdTomato negative cells (TOM$^{negwide}$) and was designed according to a tdTomato positive control sample to capture all cells exhibiting full or partial silencing memory. Scatterplots: show significant hits from the screen displayed by -log relative ranking algorithm (RRA) score from the cells captured using the TOM$^{negwide}$ gate at short-term timepoint (3d DOX-washout; left) or longer term (7d DOX-washout; right). These hits are linked with enabling epigenetic inheritance when abrogated. (**C**) Line plot for gRNA count from the CRISPR screen after silencing (+DOX) and during memory phase (DOX-washout for 3d or 7d) for the candidate genes indicated. Each line represents a different gRNA from the pool for the same target gene. A concordant enrichment during DOX washout indicates all gRNAs (and therefore independent knockouts of the target gene) promote epigenetic inheritance. Note *Kmt2d* is only enriched at the earlier timepoint.

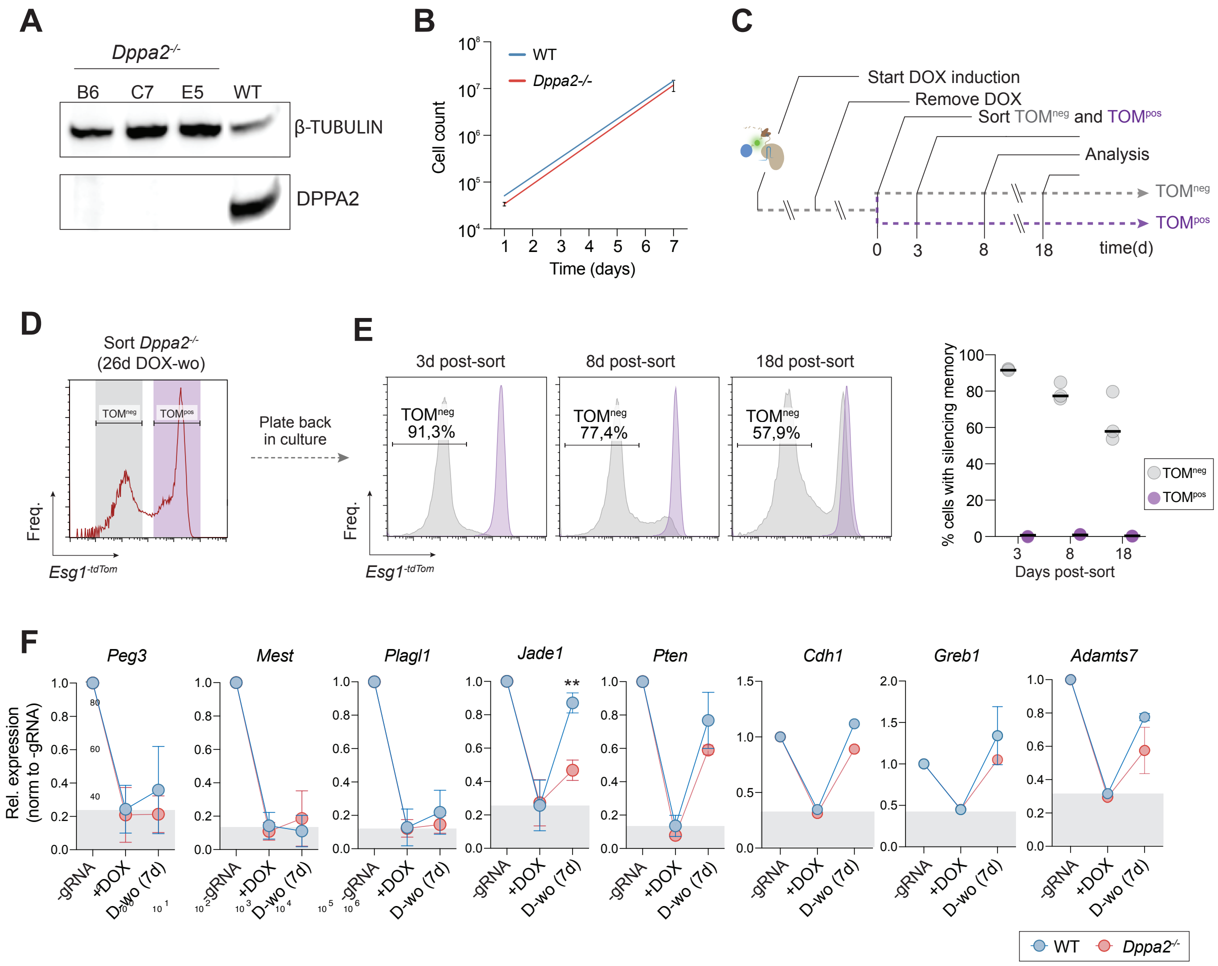


**Fig. S5. Epigenetic inheritance is probabilistic upon *Dppa2* knockout.**

(**A**) Western blot comparing DPPA2 protein expression in WT and *Dppa2* knockout individual ESC clonal lines. β-TUBULIN is used as loading control. (**B**) Line plot indicates exponential growth of wildtype or *Dppa2*$^{-/-}$ cells. (**C**) Timeline of the experiment. After induction for 7 days, tdTomato negative (TOM$^{neg}$) and positive (TOM$^{pos}$) fraction of cells are sorted following 26 days of DOX washout in *Dppa2*$^{-/-}$ and put back in culture separately. Flow cytometry analysis is performed after 3-, 8- or 18-days after sorting. (**D**) *Esg1*$^{-tdTomato}$ expression in *Dppa2*$^{-/-}$ after 26 days of DOX washout. Grey and purple boxes indicate the gates used to sort TOM$^{neg}$ and TOM$^{pos}$ fraction of cells respectively. (**E**) Left, distributions of the TOM$^{neg}$ and TOM$^{pos}$ cell population over time, showing a probabilistic reversion to active status that is cumulative, indicating a stochastic memory function. Right, percentage of silenced cells of TOM$^{neg}$ and TOM$^{pos}$ in three independent *Dppa2*$^{-/-}$ clones, after 3-, 8- or 18-days post-sort. (**F**) Line-plots show gene repression (+DOX) and memory (D-wo) of the indicated targets relative to the housekeeping gene *Rplp0* and normalised to -gRNA control. All targets exhibit increased memory in *Dppa2* kncokout, albeit insignificant, which likely reflects stochastic memory function at the single-cell level (i.e. >50% the cells still revert in *Dppa2-/-,* thereby blunting the effect in population measurements). Grey boxes represent the level of silencing achieved after +DOX treatment in the wildtype condition. Error bars are standard deviation out of two or three biological replicates. Statistics is measured by one-tail unpaired t-test between WT and *Dppa2* knockout conditions (*$**=p<0.01$*).

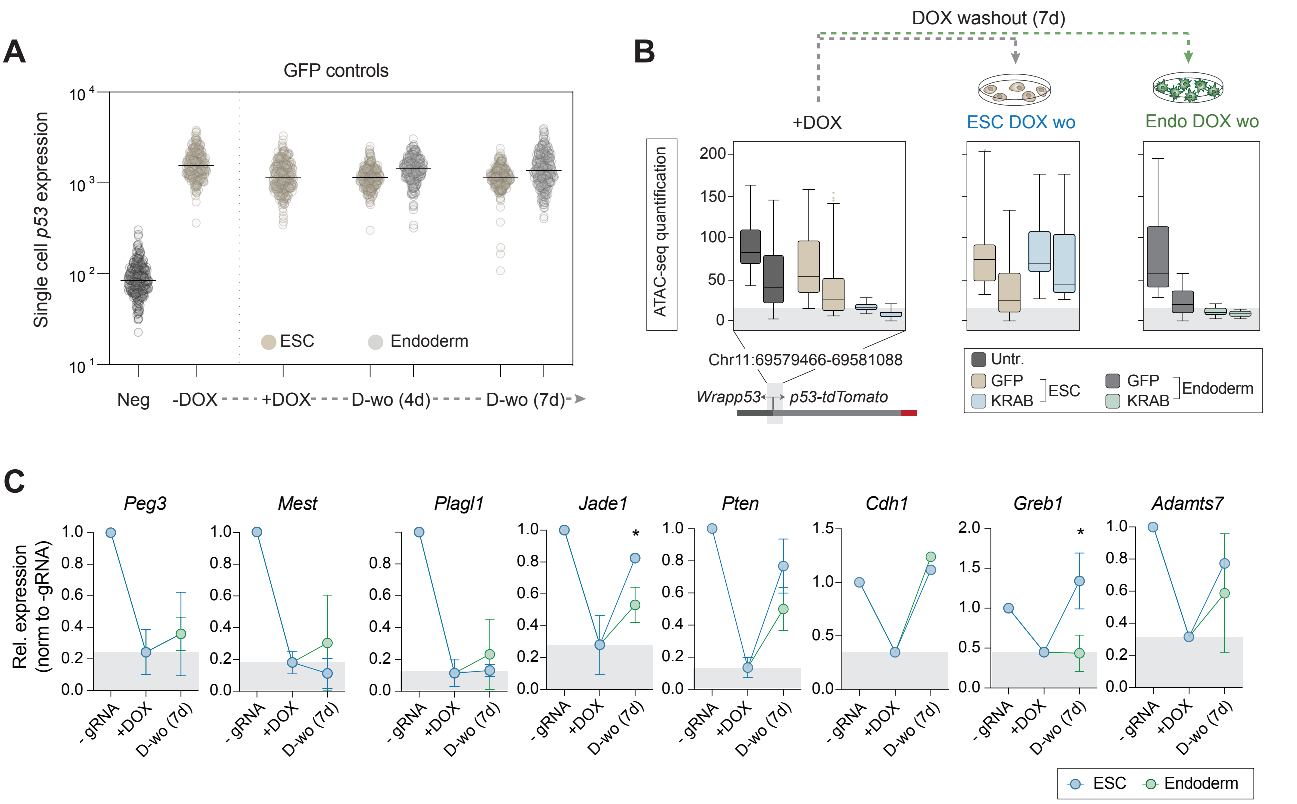


**Fig. S6. Context dependent epigenetic memory in endoderm cells**

(**A**) Expression of the *p53*$^{\text{-tdTomato}}$ reporter at the single cell level in ESC or endoderm differentiated cells in the DOX condition indicated, using only the GFP control, confirming no effect (compare to main Fig 5B). Black horizontal bars represent the median of fluorescence intensity in the population of cells. (**B**) Box plot quantification of the ATAC-seq accesibility at the extended *p53* promoter region in biological replicates. (**C**) Line-plots show expression of the indicated targets relative to the housekeeping gene *Rplp0* and normalised to -gRNA control for each timepoint in ESC or endoderm cells. They demonstrate most loci do not exhibit memory of epigenetic silencing in ESC or endoderm, but some (e.g. Jade1 and Greb1), show selective epigenetic inheritance in endoderm. Grey boxes represent the level of repression achieved after DOX treatment in the wildtype condition. Error bars are standard deviation out of two or three biological replicates. Statistics is measured by one-tail unpaired t-test between WT and knockout conditions *(*=p<0.05)*.

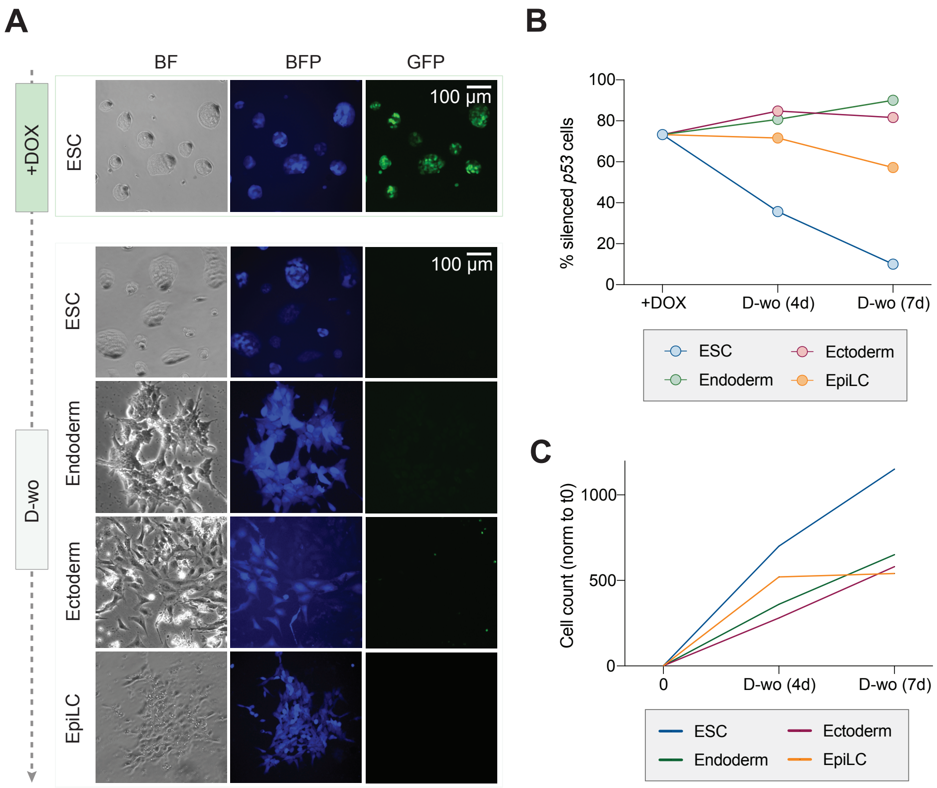


**Fig. S7 Epigenetic memory is maintained over differentiation programs representative of the three germ layers.**
(**A**) Representative microscopy images of bright field and BFP or GFP fluorescence of ESC, endoderm, ectoderm or EpiLC differentiated cells upon DOX induction (7 days) or DOX washout (7 days). (**B**) Time-course showing epigenetic memory of *p53*$^{\text{-}tdTomato}$ silencing during DOX washout in ESC, endoderm, ectoderm and EpiLC. (**C**) Line plots shows ratio of cell count normalized to timepoint 0 in ESC, endoderm, ectoderm or EpiLC differentiated cells.
